# Machine Learning-Based Drug Response Prediction Identifies Novel Therapeutic Candidates for Colorectal Cancer Cell Line KM-12

**DOI:** 10.1101/2025.02.24.639035

**Authors:** Cyrine Chanchabi, Conception Paul, Vincent Coulon, Ilaria Cicalini, Damiana Pierogostino, Sachin Vishwakarma, Saiveth Hernández-Hernández, Pedro J. Ballester, Michael Hahne

**Author notes:** shared last authorship (Pedro J. Ballester, Michael Hahne). corresponding authors: Pedro Ballester and Michael Hahne.

## Abstract

One major hurdle in the discovery of new drugs is the limited capacity of traditional screening methods to efficiently explore the immense space of drug candidates. To bypass this difficulty, computational approaches such as virtual screening (VS) have been developed supported by Machine Learning (ML) models to predict the activities of drug-like molecules on a given target. We developed a ML model for VS of a library of 25 million highly-diverse synthesis-on-demand molecules against the KM-12 colorectal cancer (CRC) cell line. By combining different biological readouts, we discovered compounds with a novel chemical scaffold, not yet described in the context of CRC, with either a prominent cytotoxic or cytostatic effect. This finding underpins the strength of our ML-guided VS protocol in discovering promising CRC drug leads triggering altered biological responses.

## Introduction

For a large number of pathologies, the development of novel therapeutics is rather long and cost intensive, as, for example, drug candidates can show important toxicity or insufficient efficacy. It is estimated that only about 1 in several thousand compounds that enter laboratory testing will ever make it to human testing (Phase 1 trials)^1^, with an average time of 12-15 years from the first tests in a research laboratory to the final drug approval and a cost of US$2–3 billion^2^.

To speed up drug development and reduce its enormous costs, computational methods have been proposed, including phenotypic VS. In this instance, ML models are developed for “drug prioritization” to reduce the search space among which the suitable compounds should be found, and select the most likely to be interesting for experimental validation. The goal is to discover as many prospective cancer drug leads as possible by predicting the activity of drug-like molecules on a specific cancer cell line. Moreover, the classical automatized drug screening approaches rely on one read-out, which might overlook specific effects of compounds.

In this study, we describe a ML approach to screen virtual libraries for novel molecules that dampen viability or proliferation of specific tumor cell lines. We chose to focus on the CRC cell line KM-12^3^, as a high number of molecules have been tested on this cell line. Importantly, in the testing of the biological responses, we employed different readouts to maximize efficiency in the selection procedure, including cell number, morphology, metabolic activity (MTS assay), cell cycle (combined BrdU incorporation/7-AAD staining), as well as cell death (i.e. apoptosis/necrosis revealed by combined Annexin V/7-AAD staining).

CRC is the second leading cause of cancer death in Europe, with >340k new cases and 150k deaths per year in the EU-27 alone, and 3 million cases per year projected worldwide (EC CRC burden Factsheet 2020)^4^. This is in spite of improved screening programs and due to the insufficient development of novel therapeutics. A new opportunity in designing novel therapeutic strategies is the recent classification of CRC into four distinct subtypes termed Consensus Molecular Subtypes (CMS): immune (CMS1), canonical (CMS2), metabolic (CMS3) and mesenchymal (CMS4)^5^. Importantly, the CMS classification correlates with responses to presently available treatments and clinical outcome. For example, CMS2 tumors respond best to existing standard-of-care (SOC) chemotherapy such as oxaliplatin, while CMS4 tumors responses to SOC are poor and are associated with the worst prognosis^6^. Thus, CMS stratification provides a robust framework for designing new subtype-specific and personalized treatments in CRC. Moreover, most of the CRC cell lines can be annotated to a specific CMS subtype therefore providing the frame for a first pre-clinical testing of subtype-specific drug responses^3^.

In this study we propose a novel ML model for VS, and we used it for the experimental identification of three novel compounds which efficiently inhibited proliferation of the CRC cell line KM-12.

## Results

### A first screen produces one promising cytotoxic compound (Dr1)

In this paper, we present a ML model for VS that aims to identify compounds targeting subtype-annotated CRC cell lines. We started by carrying out a virtual screen against the CMS1- annotated CRC cell line KM-12^3^. To this end, we developed a multi-task learning model for VS integrating the gene expression profiles of the cell lines, the chemical properties of molecules and their measured activities. This model was trained on the full NCI-60 dataset ^7^ using the XGBoost algorithm. Employing this model, we assessed over 25 million chemically-diverse compounds and selected 74 synthesis-on-demand molecules with potential KM-12 activity to test (**Screen 1: Supplementary Table 1)**. For this, KM-12 cells were treated with 2.5 micromolar (µM) of each compound for 72 hours (h). We employed the commonly used sulforhodamine B (SRB) assay for drug screening, to assess cell proliferation by measuring cellular protein content^8^. Only one of the 74 compounds (labeled Dr1) showed an important effect, with an inhibition of over 50% of cell proliferation (**Fig. 1a**). Assessment of cell viability by measuring metabolic activity using the MTS assay^9^ led to a dose-response curve similar to the SRB assay supporting the proliferation-limiting effect of Dr1 on KM-12 cells (**Fig. 1b**). However, we noted variability in the effect of the compound among biological replicates, with an IC50 (Inhibitory Concentration 50 defined as the concentration that inhibits 50% of cell proliferation) ranging between 1.5 µM and 3 µM and a standard deviation (SD) that was very important at high concentrations (**Fig. 1c**).

**Figure 1.**
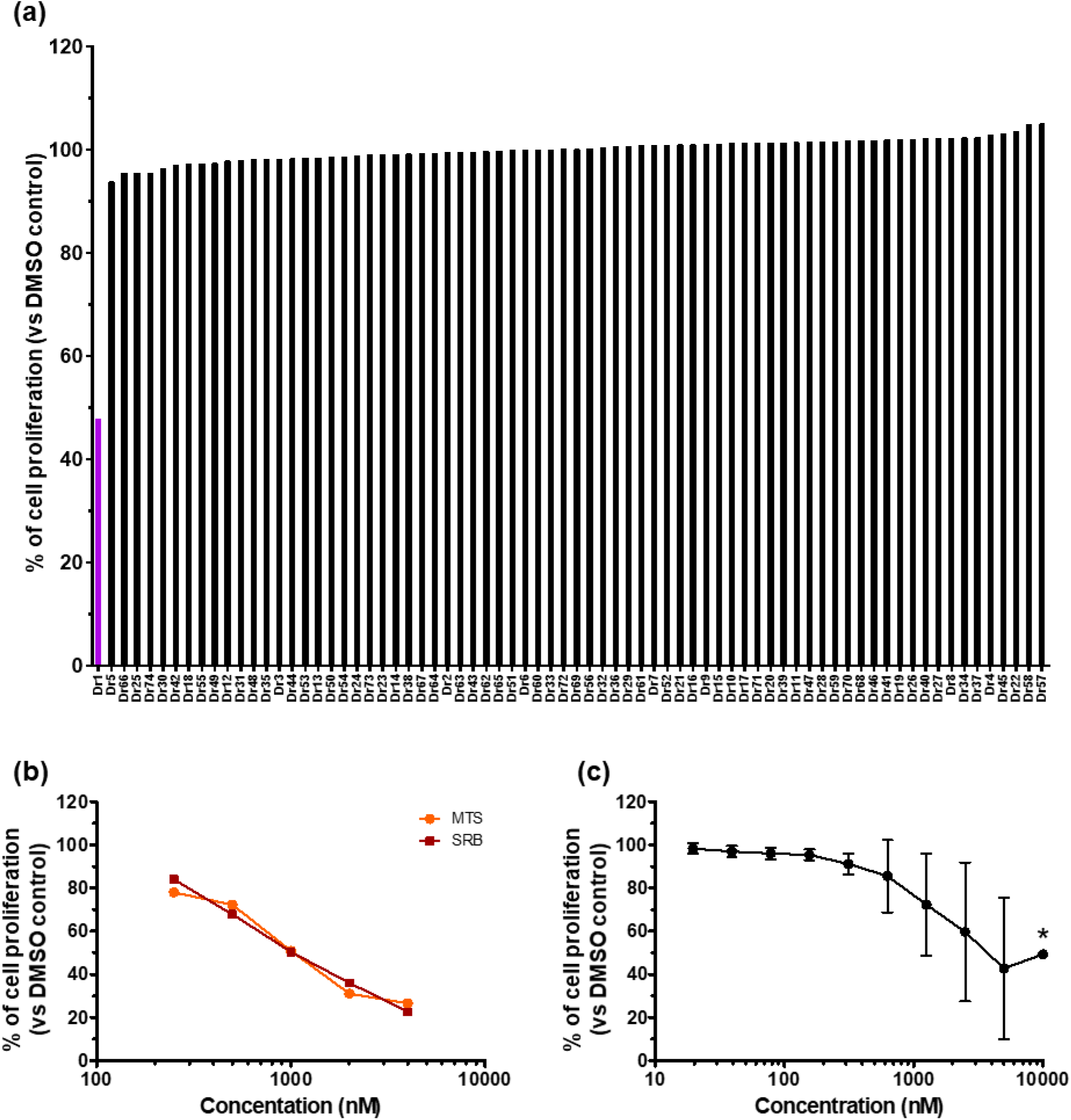
Virtual screen 1. Testing of 74 synthesis-on-demand compounds on KM-12 cells. **(a)** KM-12 cells were treated for 72 h with 2.5 µM of each designated compound. Protein content as measured by sulforhodamine B assay (SRB assay) was taken as a readout for cell proliferation. Dr1 is the only compound with an important inhibition of KM-12 cell proliferation. In purple: Most promising drug lead. KM-12 cells were cultured for 72 h at presence of the indicated concentration of the respective compound. Means of triplicates are shown. Dose-response curves of Dr1 measured by **(b)** MTS and **(c)** SRB assays, n=2. Mean +/-SD, *: n=1. Equivalent concentration of DMSO was used as control.

### A second screen generates two derivatives (Dr1-14 and Dr1-17) with improved activity

The results of this first screen were then added to the original dataset and all used to retrain the multi-task learning model to further improve KM-12 activity prediction. Next, the most similar molecules to Dr1 in the GalaXi virtual library, spanning 2.3 billion synthesis-on-demand molecules, were identified and scored with the re-trained multi-task learning model. The most promising Dr1 analogues according to this ranking were tested *in vitro*. This process resulted in the synthesis and evaluation of 39 derivatives of Dr1 (**Screen 2: Supplementary Table 2**). Employing the 2.5 µM screen described above, the two most promising derivatives (Dr1-14, Dr1-17) were selected for further testing (**Fig. 2a**). Dr1-14 was the compound with strongest cytotoxicity, particularly at high concentrations (i.e. 5 µM and 10 µM), and had an IC50 of 1.8 µM **(Fig. 2b)**. Dr1-17 was particularly effective at concentrations over 2.5 µM with a more progressive decrease of cell proliferation and a higher IC50 of 6.5 µM **(Fig. 2c)**. In contrast to Dr1, both compounds, i.e. Dr1-14 and Dr1-17, showed a reproducible effect throughout different assays with low SD.

**Figure 2.**
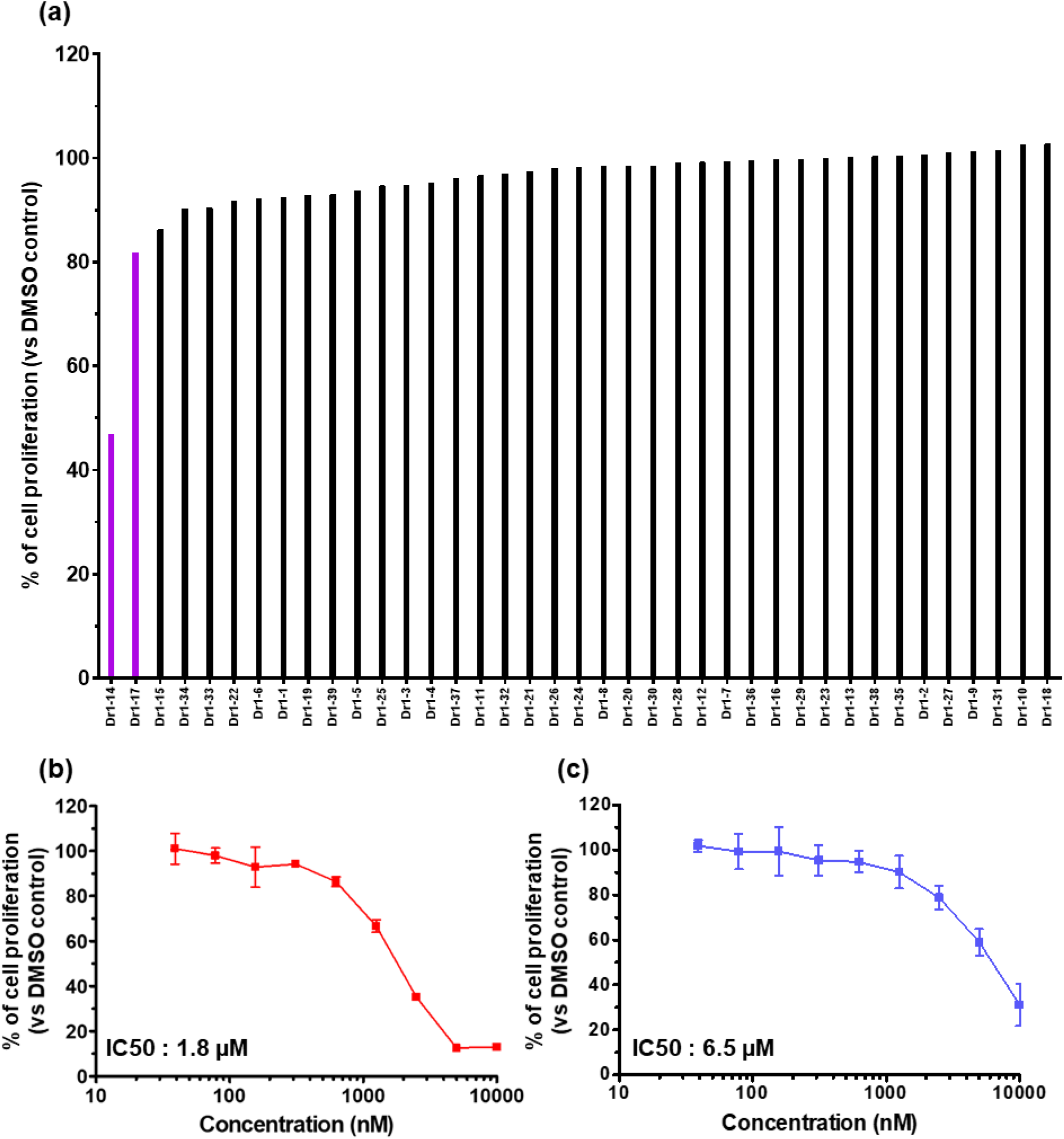
Virtual screen 2. Testing of 39 synthesis-on-demand Dr1 derivatives on KM-12 cells. **(a)** KM-12 cells were treated for 72 h with 2.5 µM of each designated compound displaying the derivatives Dr1-14 and Dr1-17 with highest proliferation inhibition activity. In purple: Most promising drug leads. KM-12 cells were cultured for 72 h at presence of the indicated concentration of **(b)** Dr1-14 and **(c)** Dr1-17 and cell proliferation was measured by SRB (sulforhodamine B) assay. n=3. Mean +/- SD. DMSO is used as control.

### Dr1-14 blocks proliferation of KM-12 cells through cell cycle arrest and cell death

We next explored the effect of Dr1-14 on KM-12 cells by treating cells with 5 or 10 µM of the drug lead or DMSO control over 72 h. The number of live cells was assessed at different time- points by trypan blue coloration. Both concentrations of Dr1-14 effectively suppressed cell proliferation already at 24 h after addition (**Fig. 3a**). This suppressed proliferation was associated with an altered morphology after 72 h by converting the cell spheroids to more isolated, rounded cells **(Fig. 3b)**. Cell-cycle analysis using 7-aminoactinomycin D (7-AAD), a fluorescent DNA intercalator, revealed that treatment of KM-12 cells with Dr1-14 led to cell cycle arrest. Indeed, 10 µM Dr1-14 significantly reduced the number of cells in G0/G1 phase as early as 5 h after treatment and led to complete disappearance of cells in G0/G1 and S phases after 24 h treatment **(Fig. 3c)**. Similar, though less pronounced, was the effect of 5 µM Dr1-14. The absence of S -phase cells after 24 h of treatment with the 10 µM concentration suggests that the remaining cells accumulate in G2/M phase (diploid cells; 4N). In addition, a proportion of the KM-12 cell population became polyploid (>4n). The disappearance of cells in G0/G1 and accumulation of cells in G2/M at 5 h suggests premature S phase entry, which should trigger DNA damage^10,11^. We therefore performed Western-blot analysis displaying elevated levels of phosphorylated H2AX (γH2AX) in Dr1-14-treated cells, indicative for DNA double-strand breaks **(Fig. 3d, Supplementary** Fig. 1). Furthermore, the increase in the subG1 fraction in Dr1-14-treated cells together with the observed DNA damage suggest that the cell arrest could lead to cell death. To verify this hypothesis, we conducted an apoptosis assay using Annexin V and 7AAD. Annexin V binds to the phosphatidyl serine (PS) exposed at the cell surface during apoptosis. 7AAD is a cell-impermeant dye that stains cells with a compromised membrane, allowing for the detection of late apoptotic and/or necrotic cells. The results show that, following the G2/M arrest, Dr1-14-treated cells undergo significant cell death through both apoptotic and necrotic pathways **(Fig. 3e, f)**.

**Figure 3.**
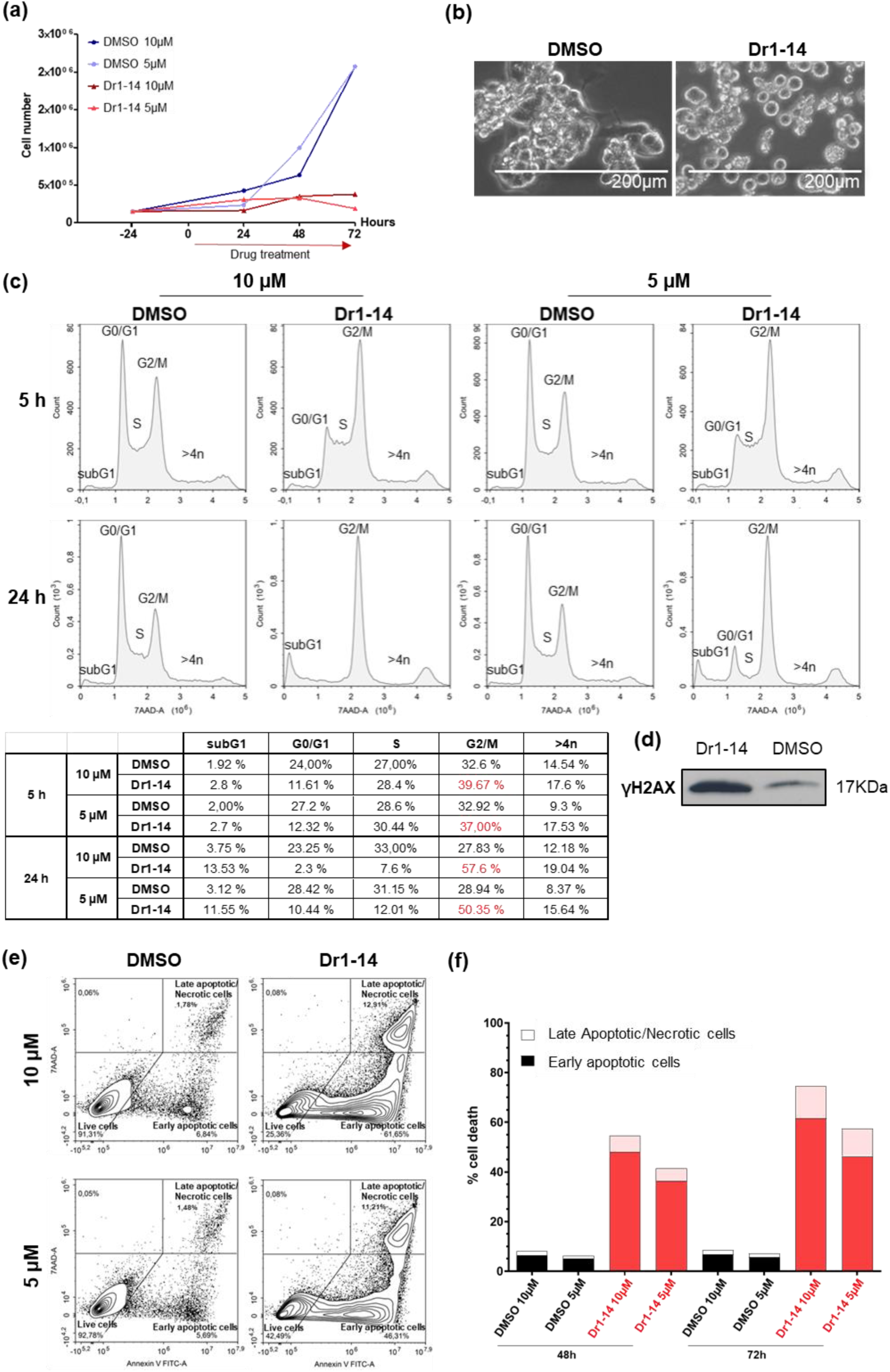
Dr1-14 induces cell cycle arrest and significant cell death in KM-12 cells. KM-12 cells were treated with 10 µM or 5 µM Dr1-14 or equivalent DMSO as a negative control. The cells were collected at the indicated time-points and submitted to various assays. **(a)** Number of live cells assessed by trypan-blue staining at 24 h, 48 h and 72 h. **(b)** Morphological changes in cells treated with 5 µM of Dr1-14 or equivalent DMSO for 72 h. **(c)** Cell cycle analysis of KM-12 cells treated with 10 µM and 5 µM of Dr1-14 or equivalent DMSO for 5 h and 24 h. The percentages of cells in each phase are reported in the table below. **(d)** Western blotting showing phosphorylated H2AX (γH2AX) levels after 24 h treatment with 5 µM of Dr1-14 or equivalent DMSO control. Loading control is shown in Supplementary Figure 1. **(e)** Gating strategy for the 72-h time-point using Annexin V and 7-AAD. Double negative: Live cells; Annexin V positive: Early apoptotic cells; Double positive: Late apop- totic and/or necrotic cells. **(f)** Percentage of early apoptotic and late apoptotic/necrotic cells after Dr1-14 or DMSO treatment for 48 h and 72 h. Each experiment was performed at least twice.

### Dr1-17 acts predominantly as a cytostatic compound

Dr1-17 had a less important effect on cell viability compared to Dr1-14 **(Fig. 4a)**, and altered differently the morphology of KM-12 cells by converting the spheroid structures into rather flattened 2D arrangements **(Fig. 4b)**. We then studied the effect of Dr1-17 on the cell cycle by flow cytometry. Using the simple 7AAD staining previously described, we observed a decrease in cells in S phase with 10 µM and 5 µM of Dr1-17 treatment in comparison to DMSO control over a 48 h time-course **(Fig. 4c)**. We next conducted a more thorough assay to study the repartition of cells in S phase by incubating Dr1-17 or DMSO-treated KM-12 cells with bromodeoxyuridine (BrdU). BrdU is a synthetic thymidine analogue that is incorporated into newly synthesized DNA, allowing to determine the percentage of cells in S phase. The BrdU-treated cells were then stained with an anti-BrdU antibody and 7-aminoactinomycine D (7- AAD) to visualize the DNA content. This combination is used to study the repartition of cells in the different phases of the cell cycle (subG1, G0/G1, S and G2/M phases)^12^. A 48 h assay revealed that Dr1-17 strongly blocks cell cycle progression at 10 µM, as the percentage of cells in S phase dropped from 35% to 14%, paralleled by an increase in the percentage of cells in G0/G1 phase **(Fig. 4d)**. The cell cycle analysis showed only a small increase of subG1 cells in response to Dr1-17 treatment. Indeed, cell death analysis using combined Annexin V/7-AAD staining displayed only low numbers of early and late apoptotic/necrotic cells **(Fig. 4e, f)**. This suggests that Dr1-17 acts predominantly as a cytostatic drug by blocking cells in G0/G1 phase, consistent with the flattened morphology.

**Figure 4.**
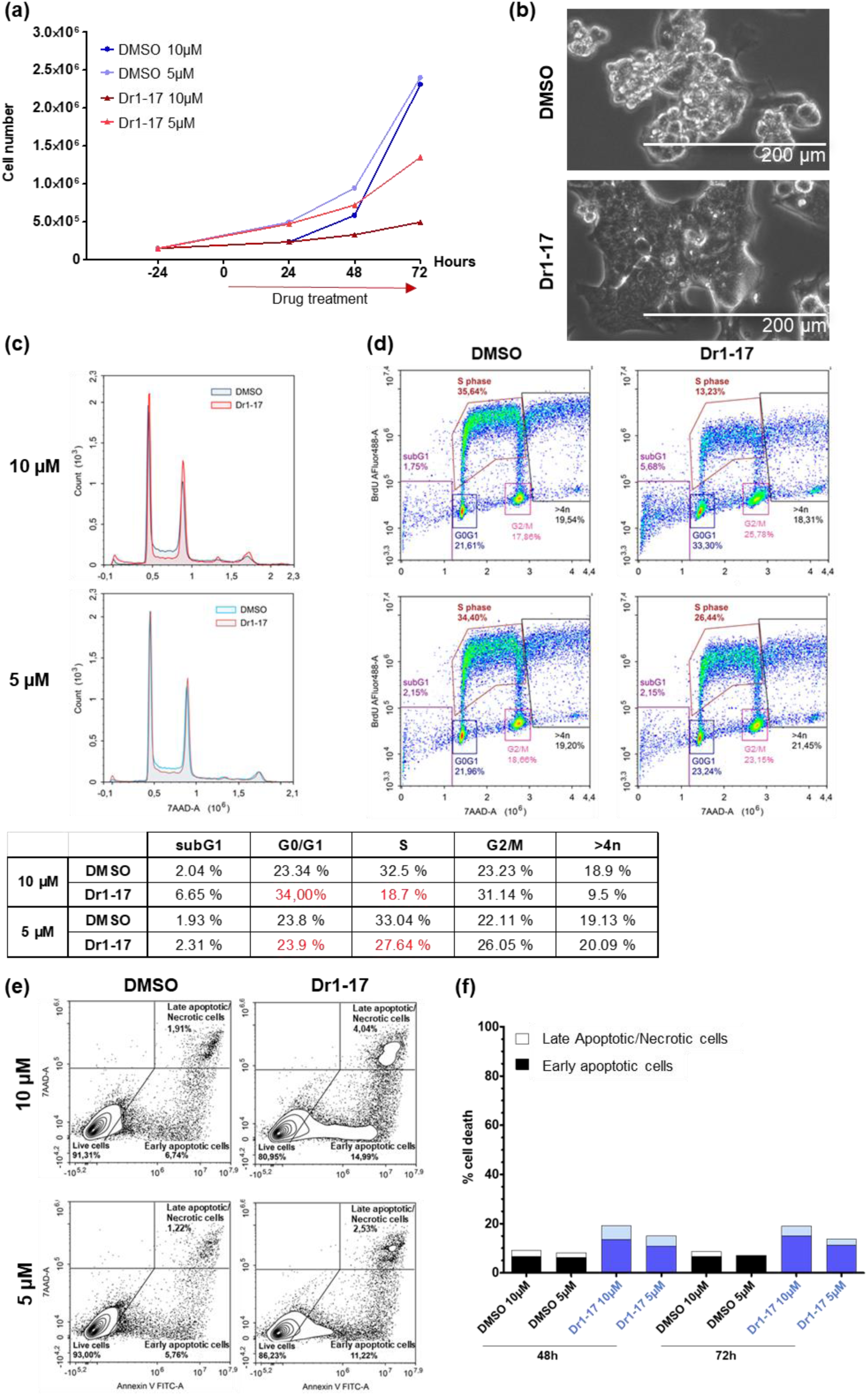
Dr1-17 induces predominantly cell cycle arrest in KM-12 cells. KM-12 cells were treated with 10 µM or 5 µM Dr1-17 or equivalent DMSO as a negative control. The cells were collected at the indicated time-points and submitted to various assays. **(a)** Number of live cells assessed by trypan-blue staining at 24 h, 48 h and 72 h. **(b)** Morphological changes in cells treated with 10 µM of Dr1-17 or equivalent DMSO for 72 h **(c)** Cell cycle analysis of KM-12 cells treated with 10 µM and 5 µM of Dr1-17 or equivalent DMSO for 48 h. The percentages of cells in each phase are reported in the table below. **(d)** BrdU-7AAD cell cycle analysis of KM-12 cells treated with 10 µM and 5 µM of Dr1-17 or equivalent DMSO for 48 h. **(e)** Gating strategy for the 72-h time-point using Annexin V and 7AAD. Double negative: Live cells; Annexin V positive: Early apoptotic cells; Double positive: Late apoptotic and/or necrotic cells. **(f)** Percentage of early apoptotic and late apoptotic/necrotic cells after Dr1-17 or DMSO treatment for 48 h and 72 h.

### A third derivative emerges from the third screen (Dr1-79)

The only difference between screen 3 and screen 2 is that virtual libraries other than GalaXi were used. Here we searched the most similar molecules to Dr1 in EnamineStore (1.2 billion compounds searched) and also in SmallWorld (4.5 billion compounds searched), to thereafter score them with the re-trained multi-task learning model. The most promising Dr1 analogues according to these rankings were tested *in vitro*. This process led to the synthesis and *in vitro* evaluation of 63 compounds (**Screen 3: Supplementary Table 3**). The 2.5 µM drug screen relying on the SRB assay revealed one particularly interesting drug lead, i.e. Dr1-79 (**Fig. 5a**), which exhibited a strong and reproducible dose-response proliferation inhibition of KM-12 cells with an IC50 of 3.2 µM (**Fig. 5b**). Although this IC50 is higher than that of Dr1-14 (1.8 µM), it is still lower than the one of Dr1-17 (6.5 µM) **(Fig. 5c)**. We next explored the mechanisms underlying the proliferation-blocking effect of Dr1-79. For this, we treated KM-12 cells with higher doses of the compound (10 and 5 µM) and collected them at different time-points to analyze cell cycle and cell death. Treatment of KM-12 cells with Dr1-79 induced similar morphological alterations as observed for Dr1-17, i.e. a more “flattened” cellular shape and less spheroid-like structures, but decreased the number of live cells more efficiently **(Fig. 6a, b)**. Accordingly, the effect of Dr1-79 on cell cycle was similar to the one observed for Dr1-17 with cells disappearing from S phase and accumulating in G0/G1. Notably, these alterations were also detectable in the 5 µM condition **(Fig. 6c, d)**. In parallel to the BrdU cell cycle analysis, we carried out an apoptotic assay using the Annexin V/7-AAD staining strategy. This analysis showed that Dr1-79 induces a modest early apoptotic (10%) and late apoptotic/necrotic (2-4%) response in KM-12 cells that can account for a part of the decrease in the number of live cells **(Fig. 6e, f)**. Together, these results indicate that Dr1-79 inhibits KM- 12 cells proliferation at high doses and promotes cell death to a certain extent.

**Figure 5.**
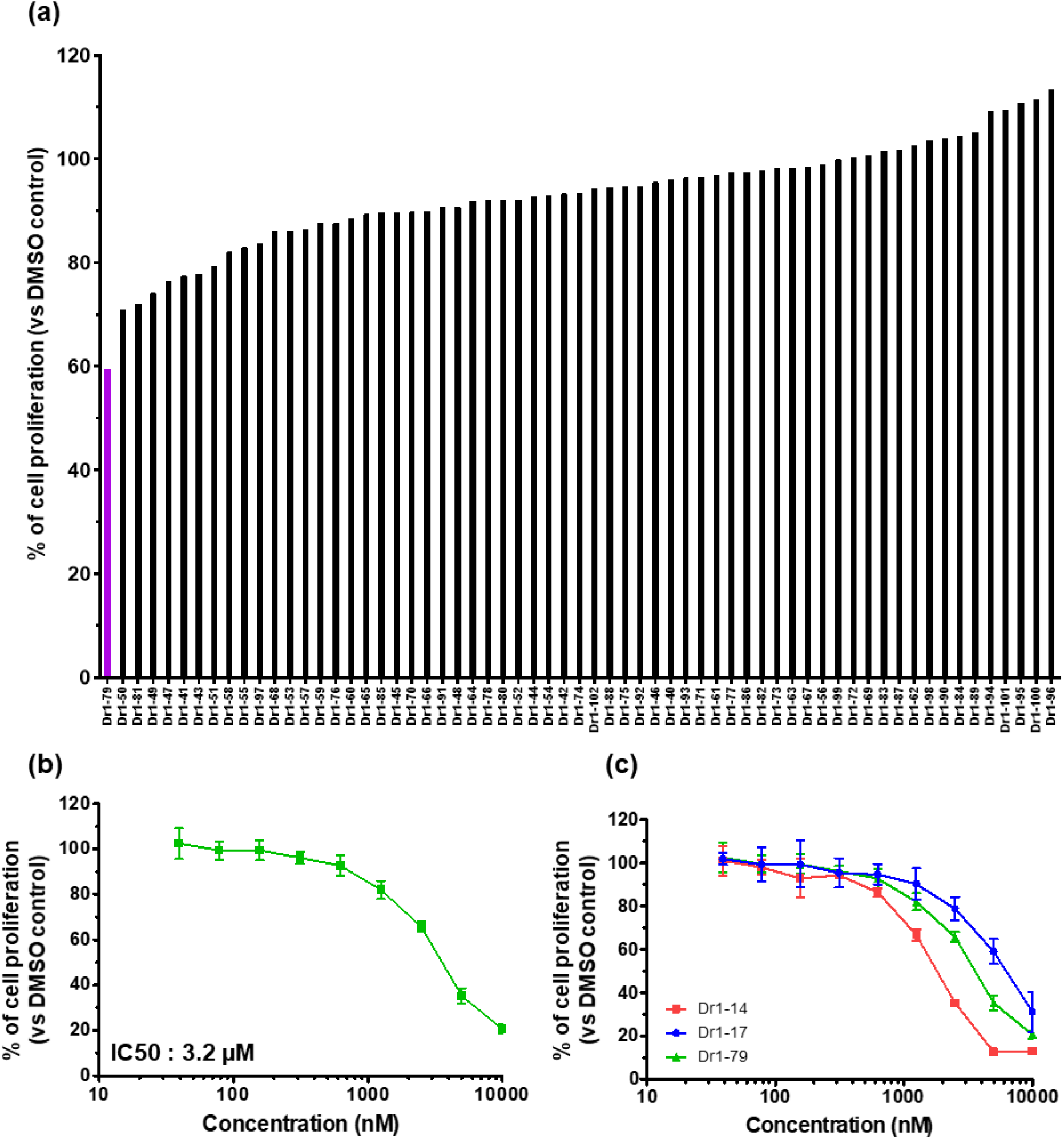
Virtual screen 3. Testing of 63 synthesis-on-demand derivatives from compounds down-selected from the second screen on KM-12 cells. **(a)** KM-12 cells were treated for 72 h with 2.5 µM of each designated compound. One derivative (Dr1-79) shows the most important cell proliferation inhibition effect on KM-12 cells. In purple: Most promising drug leads. **(b)** KM-12 cells were cultured for 72 h at presence of the indicated concentration of Dr1-79 and the dose-response curve was measured by sulforhodamine B assay, n=3 **(c)** Comparison of dose-response inhibition of the three main drug leads we identified: Dr1-14, Dr1-17, Dr1-79. n=3. Mean +/- S. DMSO is used as control.

**Figure 6.**
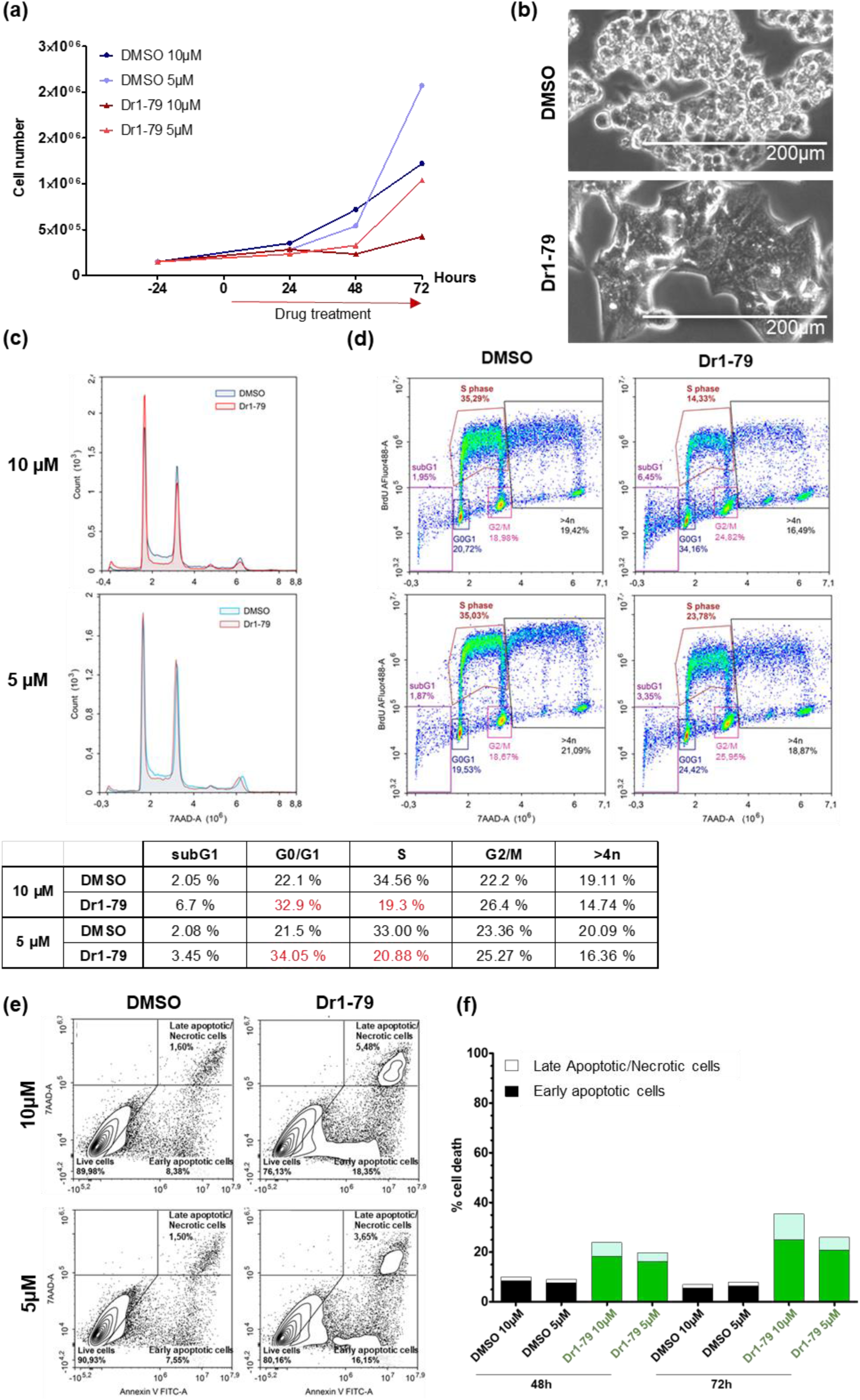
Dr1-79 induces cell proliferation arrest and moderate cell death in KM-12 cells. KM-12 cells were treated with 10 µM or 5 µM of Dr1-79 or equivalent DMSO as a negative control. The cells were collected at the indicated time-points and submitted to various assays. **(a)** Number of live cells assessed by trypan-blue staining at 48 h and 72 h. **(b)** Morphological changes in cells treated with 10 µM of Dr1-79 or equivalent DMSO for 48 h. **(c)** Cell cycle analysis of KM-12 cells treated with 10 µM and 5 µM of Dr1-79 or equivalent DMSO for 48 h. The percentages of cells in each phase are reported in the table below. **(d)** BrdU/7-AAD cell cycle analysis of KM-12 cells treated with 10 µM and 5 µM of Dr1-79 or equivalent DMSO for 48 h **(e)** Gating strategy for the 72-h time-point using Annexin V and 7-AAD. Double negative: Live cells; Annexin V positive: Early apoptotic cells; Double positive: Late apoptotic and/or necrotic cells. **(f)** Percentage of early apoptotic and late

As Dr1-79 caused a decrease in cell number and inhibited proliferation but induced low to moderate cell death, we next sought to determine if this was representative of a reversible or non-reversible response. To explore this hypothesis, KM-12 cells treated with 10 µM of Dr1-79 for 72 h were released in normal medium (without drug lead) for an additional 72 h. The cell-cycle analysis shows that the cells start proliferating again after the release, with percentages of cells in each phase of the cell cycle that are similar to the 72 h-treated DMSO control **(Fig. 7a)**. The morphology of the Dr1-79-treated cells also goes back to normal, similar to the DMSO control **(Fig. 7b)**. Thus, Dr1-79 acts predominantly as a cytostatic agent, as previously defined^13^, that displays a transient inhibitory effect on KM-12 cell proliferation,.

**Figure 7.**
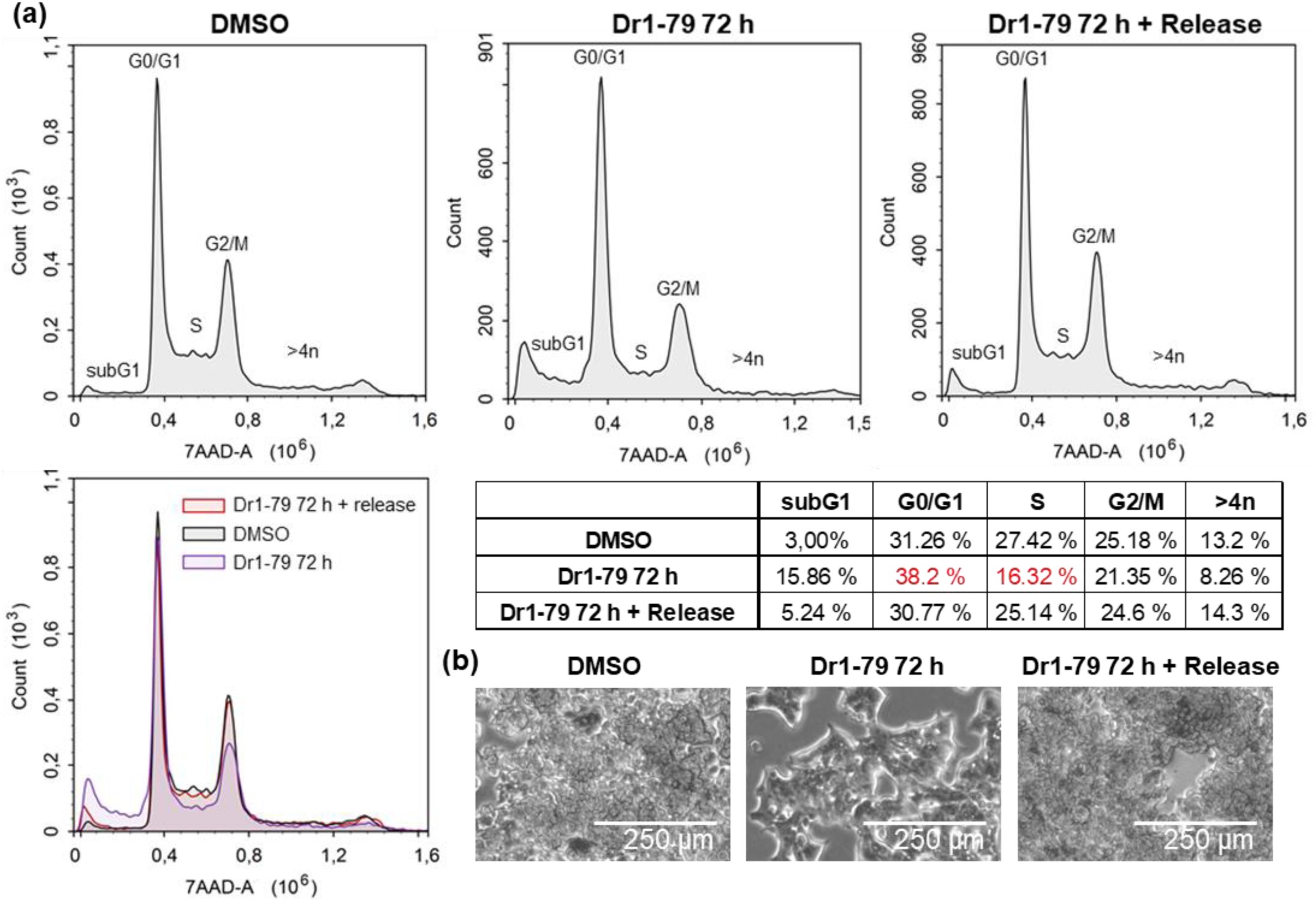
The effect of Dr1-79 on KM-12 cells is reversible. KM-12 cells were treated with 10 µM of Dr1-79 or equivalent DMSO control for 72 h. The cells were then released in normal medium without treatment for an additional 72 h and collected to analyze the cell cycle by flow cytometry. **(a)** Cell cycle analysis. The percentages of cells in each phase are reported in the table below **(b)** Morphological changes in KM-12 cells after 72 h of treatment with Dr1-79 or DMSO control and after release in normal medium.

## Discussion

The importance of ML for VS has not stopped growing in recent years. This has been based on the availability of large virtual libraries and chemical spaces of compounds that can be synthesized from available building blocks with high probability and in short time ^14,15,16^. The most developed branch is ML for structure-based VS^17–19^. Indeed, VS has been successfully employed in docking experiments ^20^. Docking refers to the computational process of predicting the optimal position, orientation, and conformation of a small molecule when bound to a specific site on a target protein. Examples for successful docking experiments in the frame of structure based VS include the identification of potent inhibitors for AmpCbeta-lactamase (Amp) and agonists for the D4 dopamine receptors ^21^, novel antibiotics ^22^, as well as senolytic drugs ^23,24^.

Here we have focused on the less explored application of machine learning to phenotypic VS, where the label to predict is the viability inhibition induced by the molecule on a given cell culture. We present a prospective application of machine learning for VS of a diverse synthesis-on-demand library, i.e. the Real database available at Enamine^25^, against CRC cell lines. More specifically, we developed a multi-task learning model for VS integrating the gene expression profiles of the cell lines, the chemical properties of molecules and their measured activities. This model was trained on the full NCI-60 dataset ^7^ using the XGBoost algorithm. Employing this model, we assessed over 25 million chemically diverse molecules and selected 74 synthesis-on-demand molecules with potential KM-12 activity. Then we tested these compounds on KM-12 cells by employing the commonly used sulforhodamine B (SRB) assay to assess cell proliferation^8^. This screen revealed one compound with a particular cell viability- dampening capacity (Dr1). The results of this first screen were then integrated in the algorithm to improve compound prediction. This loop of VS and biological response testing was repeated two additional times. Notably, we observed an increase in the percentage of compounds reducing cell viability below 90% from screen 1 to 3 confirming efficacy of the applied machine learning approach **(Supplementary Table 4)**, from 1.4% of compounds decreasing the viability below 90% in the first screen, to 35% of them in screen 3 **(Fig. 9)**.

Three promising drug leads emerged from the 176 compounds tested in total. These leads labeled Dr1-14, Dr1-17 and Dr1-79 efficiently inhibited KM-12 cell proliferation, showing a reproducible dose-response inhibitory effect on the CRC cell line **(Fig. 5c**, **Fig. 8)**. The compound identified in the initial screen, i.e. Dr1, showed some variation in efficiency between different assays. This was not the case for the aforementioned molecules, suggesting that they have an improved stability.

**Figure 8.**
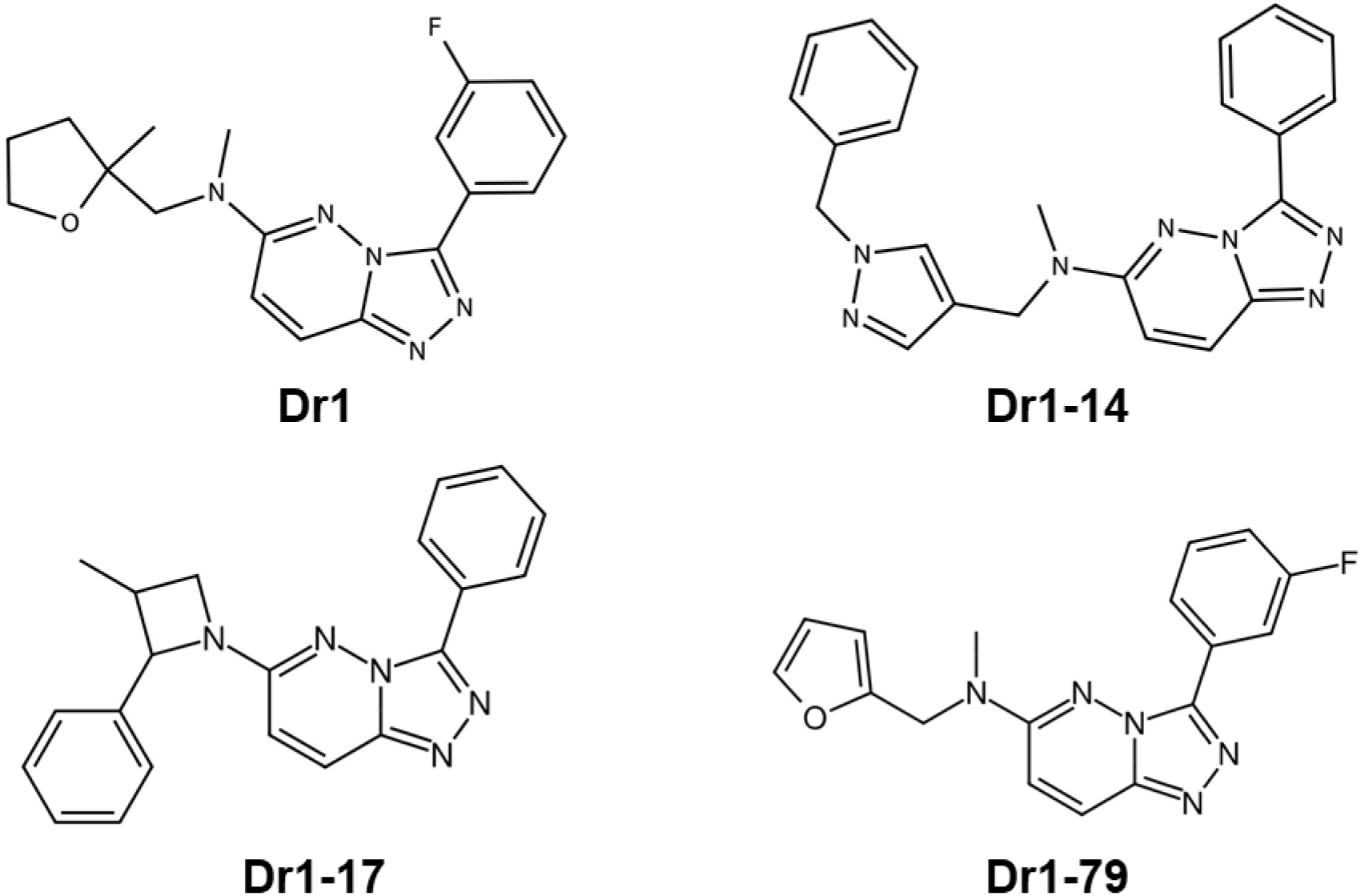
Chemical structure of the drug leads selected in virtual screens 1-3. The four compounds all share the same [1,2,4]triazolo[4,3-b]pyridazin core.

**Figure 9.**
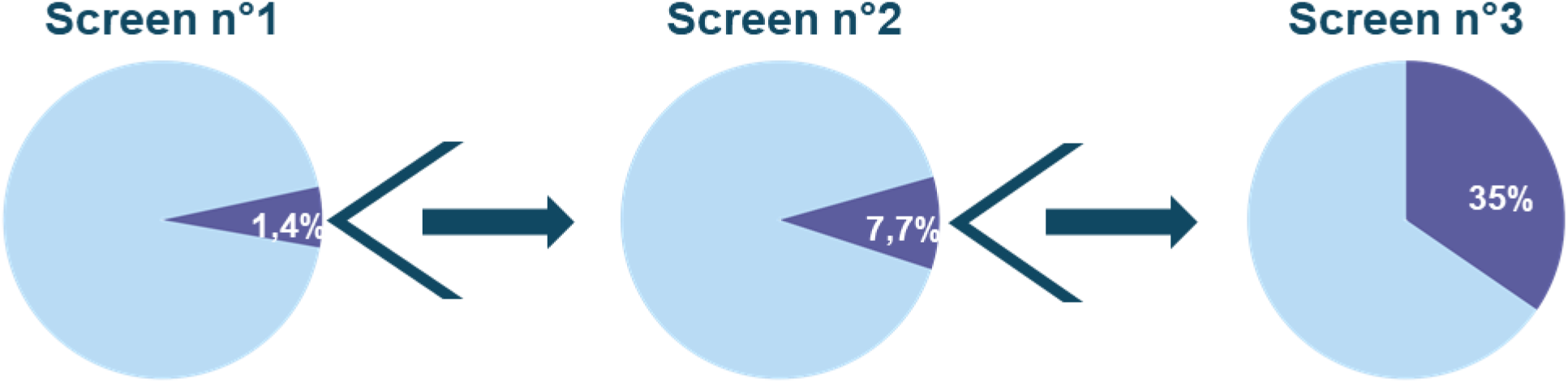
Increased success rate of identifying novel compounds that decrease cell proliferation below 90% in a 72 h SRB assay (at 2.5 µM).

Our initial intention has been to identify novel compounds of low IC50 as frequently used to monitor drug efficiency. We noticed, however, that compounds with a relatively high IC50 efficiently induced important morphological alterations and decreased cell proliferation at elevated concentrations (5-10 µM). It has been reported that the IC50 does not necessarily reflect the efficacy of cytostatic drugs^26^. Notably, a large number of reported IC50 are higher than the actual maximal concentration tested^27^. We therefore decided to use additional readouts to monitor compound efficiency such as morphological alterations, metabolic activity, cell cycle and cell death. Combining these different biological readouts to monitor compound activity identified the above-mentioned molecules, Dr1-14, Dr1-17, Dr1-79, with specific effects on cell morphology, cell death and cell cycle, respectively, all decreasing cell proliferation down to 20% in a three days treatment. Importantly, the compounds contain a similar molecular scaffold **(Fig. 7)**, which is distinct from previously described tumor growth modulating drugs. Notably, in spite of a common core structure, the down-selected compounds demonstrate a different mode of action, with Dr1-14 being cytotoxic, whereas Dr1-17 and Dr1-79 display important cytostatic features. The cell cycle analysis suggests that Dr1-17 and Dr1-79 block the transition from G0/G1 to S phase, thus allowing cells to survive^28^. Despite the fact that these two compounds act in a similar fashion, Dr1-79 has a lower IC50 (i.e. 3.2 µM) than Dr1- 17 (i.e. 6.5 µM), inhibits proliferation more strongly at lower doses (5 µM) and induces more cell death. These differences are in favor of an improvement in the efficacy of the cytostatic compounds thanks to the machine learning approach as Dr1-79 belongs to the latest screen.

While this manuscript was in preparation He and colleagues reported an approach complementary to ours for identifying novel compounds dampening viability of CRC cells^29^. This elegant work combined neural networks trained on compounds reported to be bioactive on CRC cells and curated in the ChEMBL database as well as prediction models to forecast novel molecules decreasing cell viability on the CRC cell line HT-29. Though distinct to our presented strategy, the report of He *et al.* promotes the concept of integrating computational approaches for the identification of novel therapeutic compounds.

In summary, in this article we provide proof-of-principle for the rationale to combine VS and machine learning approaches for identifying novel compounds for targeting specific tumor types. This will overcome hurdles of classical automated drug-screening approaches, which are limited by the number of compounds that can be tested. Our functional approach focuses on the impact on cell fate instead of a single target protein, allowing the identification of useful candidate molecules whatever the target. We expect that ongoing work will develop improved algorithms for compound prediction. In addition, future studies will reveal the molecular targets of the selected compounds, and thus additional vulnerabilities for targeting KM-12 cells and related tumors.

## Material and Methods

### Screening libraries

The diverse set of Enamine REAL, containing 25,270,743 molecules, was employed for screen 1 leading to the discovery of Dr1. The most similar molecules to Dr1 in the GalaXi virtual library, spanning 2.3 billion synthesis-on-demand molecules, were searched in screen 2. This is WuXi AppTec Virtual Space^30^, which performed this similarity search for us using a 2048-BIT ECFP4 fingerprint. In screen 3, the most similar molecules to Dr1 in EnamineStore (https://enaminestore.com/search) among its 1.2 billion compounds and also in SmallWorld (https://sw.docking.org/search.html) among its 4.5 billion compounds were identified.

### Machine learning modelling

We considered 2,707,434 labelled instances from pairing 50,156 unique molecules with SMILES and 60 cancer cell lines with their gene expression profiles from the NCI-60 repository (https://dtp.cancer.gov/discovery_development/nci-60/). The label is pGI50, the negative logarithm of the half-maximal growth inhibition concentration of the molecule in molar units when tested on the cell line^31^. Each molecule is featurised using 256-bit Morgan fingerprints with radius 2 and each cell line is featurised using Spearmann rank-correlation kernel on the gene expression profiles^32^. In addition to using Morgan fingerprints, we enriched our feature set by including several physicochemical properties of the compounds. These properties include total polar surface area^33^, molecular weight^34^, LogP (Partition coefficient)^35^, number of aliphatic rings, number of aromatic rings^36^, number of hydrogen bond acceptors, and number of hydrogen bond donors^37^. A XGBoost algorithm was employed to build this multi-task model using the hyperparameter values from a related drug synergy project^38^. The XGBoost model was selected due to its demonstrated strong performance on the NCI-60 dataset, effectively managing its complexity and yielding robust results^39,40^. This model was used to perform screen 1 and retrained with the addition of screen 1 results for screens 2 and 3.

### Cell culture

KM-12 cells were kindly provided by Dr. Jan Paul Medema’s lab (AMC, Amsterdam, The Netherlands; authenticated by STR sequencing). The cell line was cultured in RPMI 1640 with L-glutamine, added 25 mM HEPES (Thermo-Fisher Scientific), 10% fetal bovine serum, penicillin and streptomycin, 1% D-glucose solution plus (Sigma-Aldrich) and 100 μM sodium pyruvate (Life Technologies). The cytotoxic compounds were resuspended in DMSO at a concentration of 2 mM (screen 1 and 2) and 5 mM (screen 3).

### Sulforhodamine B assay

Twenty-four hours prior to their treatment with a drug lead or DMSO (negative control), KM-12 cells were seeded in 96-well plates at a density of 5 x 10^3^ cells per well. After 3 days of treatment, sulforhodamine B (SRB) assays were performed according to previously described procedures^8^. % cell proliferation = (mean OD510 of treated cells / mean OD510nm of control cells) x 100. Notably, this assay is based on the measurement of cellular protein content and only provides an indirect readout for cell proliferation.

### MTS assay

KM-12 cells were seeded in 96-well plates at a density of 5 x 10^3^ cells/well. After 24 hours, the cells were incubated with various concentrations of Dr1 (from 4 up to 0.18 µM) or equivalent DMSO as a negative control. Three days later, 30 µL of the MTS reagent (1 :20 parts of Phenazine Methosulfate PMS (0.92 mg/ml in PBS) and MTS (2 mg/ml in PBS), Promega) was added to each well and the plates were incubated for an additional 4 h at 37°C. The optical density was recorded at 490 nm. This assay measures metabolic activity which is also an indirect readout for cell proliferation.

### Trypan blue-based cell counting

KM-12 cells were seeded in 6-well plates at a density of 1.5 x 10^5^ cells/well and left to adhere for 24 hours. The cells were then treated with 10 or 5 µM of drug lead or equivalent DMSO concentration as negative control. They were collected by trypsinization at different time- points, along with the supernatant and the washing PBS. The total cell suspension was counted using 0.4 % trypan blue (Invitrogen, Thermofisher Scientific) staining in an automated Countess III cell counter.

### Apoptosis assay

KM-12 cells were seeded in 6-well plates at a density of 1.5 x 10^5^ cells/well. After 24 hours, they were treated with 10 or 5 µM of drug lead or equivalent DMSO as a negative control. The adherent and floating cells were collected with trypsin at different time-points and washed once in Annexin V staining buffer (0.1 M Hepes pH=7.4, 1.4 M NaCl, 25 mM CaCl2). They were then incubated with 5 µL of FITC conjugated Annexin V (Biolegend) and 5 µL of 7-AAD viability staining solution (Biolegend) in 100 µL of Annexin V staining buffer, according to the manufacturer’s guidelines. After a 15-minute incubation at room temperature, the results were acquired on a NovoCyte®Flow Cytometer (ACEA Biosciences).

### Cell cycle (DNA content) assay

KM-12 cells were seeded in 6-well plates at a density of 1.5 x 10^5^ cells/well and left to adhere for 24 hours. The cells were then treated with 10 or 5 µM of Dr1-14 or equivalent DMSO concentration as negative control. The samples were dissociated with trypsin at different time- points (5 h, 24 h), counted with trypan blue and washed twice with PBS. Ice-cold 70° ethanol was used to fix the cells (overnight to several days). The fixed cells were washed twice with PBS and stained with 7-AAD (25 µg/mL in PBS) for 30 minutes at room temperature. The results were acquired on a NovoCyte®Flow Cytometer (ACEA Biosciences).

### Bivariate BrdU / 7-AAD cell cycle analysis

KM-12 cells were seeded in 6-well plates at a density of 1.5 x 10^5^ cells/well. Twenty-four hours later, they were treated with the compound (10 µM, 5 µM) or equivalent DMSO as a negative control. After 48 h, the cells were pulsed with 100 µM of Bromodeoxyuridine (BrdU, 100 mM in DMSO) for 1 hour at 37°C then collected by trypsinization and counted with trypan blue. The labeled cells were washed once with cold PBS and fixed in ice-cold 90° ethanol (overnight to several days). Prior to the staining, they were washed twice with cold PBS then DNA was denatured for 30 minutes in 1 ml of a solution of 2N hydrochloric acid (HCl)-0.5% triton X-100. Following denaturation, the acid was neutralized with 1 ml of sodium tetraborate buffer (Na2B4O7, pH=8.5). The cells were then centrifuged and incubated with 5µL of Alexa Fluor 488-conjugated anti-BrdU antibody (3D4 clone, Biolegend) diluted in FACS buffer (PBS-0.5% Tween 20-1% BSA) for 1 hour at room temperature. After one last wash step in PBS, the cells were incubated with 7-AAD staining solution (0.14 µg/ml 7-AAD in PBS) for 30 minutes at room temperature. The results were acquired on a NovoCyte®Flow Cytometer (ACEA Biosciences).

### Western blotting

KM-12 cells were seeded in 6-well plates at a density of 1.5 x 10^5^ cells/well. After 24 hours, they were treated with 5 µM of Dr1-14 or equivalent DMSO as a negative control. Twenty-four hours post-treatment, the plates were placed on ice, washed with cold PBS and scraped with a cell scraper. The collected cells were centrifuged for 5 min at 300g at 4°C and then washed once more with cold PBS. The supernatant was removed, the pellets were frozen in liquid nitrogen and kept at -80°C until analysis. To extract the proteins, the cells were incubated in lysis buffer (50 nM Tris-HCl, pH=8; 150 mM NaCl; 1% SDS; 1% Triton X-100; Complete protease inhibitors (Roche); PhoStop phosphatase inhibitors (Roche)) for 20 minutes on ice followed by a sonication step. The protein concentration was measured using a BCA assay following the manufacturer’s instructions (Pierce™ BCA Protein Assay Kit, Thermoscientific). 10 µg of proteins were loaded on a 4-15% Mini-PROTEAN TGX SDS-PAGE Gel (Biorad). After migration in Tris-glycine buffer, the proteins were transferred on a nitrocellulose membrane using a semi-dry transfer method on the Bio-Rad Trans-Blot Turbo Transfer System according to the manufacturer’s instructions (Biorad). γH2AX was revealed using a primary antibody (Rabbit polyclonal, anti-phospho Ser139 γH2AX, 1:5000, abcam) incubated 1 h at 4°C. A secondary goat anti-rabbit HRP antibody was used to reveal the primary (1:10000, 30 minutes). The final revelation was done on the Chemidoc MP Imaging System (Biorad).

## Acknowledgments

We are grateful to the Montpellier Ressources Imagerie platform. Many thanks to Thierry Gostan and the data analysis facility SERANAD for help on the statistical analysis, and the Montpellier Ressources Imagerie (MRI) platform. This work was supported by Cancer Inserm- MIC 2021. Authors are thankful to WuXi AppTec for their support and D. Raimondi for reading carefully the article.

## References

1. Sun, D., Gao, W., Hu, H. & Zhou, S. Why 90% of clinical drug development fails and how to improve it? Acta Pharm. Sin. B 12, 3049–3062 (2022).

2. Singh, N. et al. Drug discovery and development: introduction to the general public and patient groups. Front. Drug Discov. 3, (2023).

3. Linnekamp, J. F. et al. Consensus molecular subtypes of colorectal cancer are recapitulated in in vitro and in vivo models. Cell Death Differ. 25, 616–633 (2018).

4. https://encr.eu/sites/default/files/inlinefiles/Colorectal_cancer_factsheet_March_2021.pdf.

5. Guinney, J. et al. The consensus molecular subtypes of colorectal cancer. Nat. Med. 21, 1350–1356 (2015).

6. Linnekamp, J. F. et al. Consensus molecular subtypes of colorectal cancer are recapitulated in in vitro and in vivo models. Cell Death Differ. 25, 616–633 (2018).

7. Shoemaker, R. H. The NCI60 human tumour cell line anticancer drug screen. Nat. Rev. Cancer 6, 813–823 (2006).

8. Vichai, V. & Kirtikara, K. Sulforhodamine B colorimetric assay for cytotoxicity screening. Nat. Protoc. 1, 1112–1116 (2006).

9. Soman, G. et al. MTS dye based colorimetric CTLL-2 cell proliferation assay for product release and stability monitoring of Interleukin-15: Assay qualification, standardization and statistical analysis. J. Immunol. Methods 348, 83–94 (2009).

10. Bartkova, J. et al. Oncogene-induced senescence is part of the tumorigenesis barrier imposed by DNA damage checkpoints. Nature 444, 633–637 (2006).

11. Teixeira, L. K. et al. Cyclin E deregulation promotes loss of specific genomic regions. Curr. Biol. CB 25, 1327–1333 (2015).

12. Bialic, M., Al Ahmad Nachar, B., Koźlak, M., Coulon, V. & Schwob, E. Measuring S-Phase Duration from Asynchronous Cells Using Dual EdU-BrdU Pulse-Chase Labeling Flow Cytometry. Genes 13, 408 (2022).

13. Gautam, P. et al. Identification of selective cytotoxic and synthetic lethal drug responses in triple negative breast cancer cells. Mol. Cancer 15, 34 (2016).

14. Sadybekov, A. V. & Katritch, V. Computational approaches streamlining drug discovery. Nature 616, 673–685 (2023).

15. Adeshina, Y. O., Deeds, E. J. & Karanicolas, J. Machine learning classification can reduce false positives in structure-based virtual screening. Proc. Natl. Acad. Sci. 117, 18477–18488 (2020).

16. Wallach, I. et al. AI is a viable alternative to high throughput screening: a 318-target study. Sci. Rep. 14, 7526 (2024).

17. Ballester, P. J. & Mitchell, J. B. O. A machine learning approach to predicting protein– ligand binding affinity with applications to molecular docking. Bioinformatics 26, 1169– 1175 (2010).

18. Ballester, P. J. et al. Hierarchical virtual screening for the discovery of new molecular scaffolds in antibacterial hit identification. J. R. Soc. Interface 9, 3196–3207 (2012).

19. Tran-Nguyen, V.-K., Junaid, M., Simeon, S. & Ballester, P. J. A practical guide to machine-learning scoring for structure-based virtual screening. Nat. Protoc. 18, 3460– 3511 (2023).

20. Zhu, H., Zhang, Y., Li, W. & Huang, N. A Comprehensive Survey of Prospective Structure-Based Virtual Screening for Early Drug Discovery in the Past Fifteen Years. Int. J. Mol. Sci. 23, 15961 (2022).

21. Lyu, J. et al. Ultra-large library docking for discovering new chemotypes. Nature 566, 224–229 (2019).

22. Stokes, J. M. et al. A Deep Learning Approach to Antibiotic Discovery. Cell 180, 688–702.e13 (2020).

23. Smer-Barreto, V. et al. Discovery of senolytics using machine learning. Nat. Commun. 14, 3445 (2023).

24. Wong, F., Omori, S., Donghia, N. M., Zheng, E. J. & Collins, J. J. Discovering small-molecule senolytics with deep neural networks. *Nat*. Aging 3, 734–750 (2023).

25. Grygorenko, O. O. et al. Generating Multibillion Chemical Space of Readily Accessible Screening Compounds. iScience 23, 101681 (2020).

26. Brooks, E. A. et al. Applicability of drug response metrics for cancer studies using biomaterials. Philos. Trans. R. Soc. Lond. B. Biol. Sci. 374, 20180226 (2019).

27. Codicè, F. et al. The Specification Game: Rethinking the Evaluation of Drug Response Prediction for Precision Oncology. Preprint at 10.1101/2024.10.01.616046 (2024).

28. Pennycook, B. R. & Barr, A. R. Restriction point regulation at the crossroads between quiescence and cell proliferation. FEBS Lett. (2020) doi:10.1002/1873-3468.13867.

29. He, D. et al. De Novo Generation and Identification of Novel Compounds with Drug Efficacy Based on Machine Learning. Adv. Sci. 11, 2307245 (2024).

30. Virtual Screening - WuXi Biology. https://wuxibiology.com/drug-discovery-services/hit-finding-and-screening-services/virtual-screening/.

31. Piyawajanusorn, C., Nguyen, L. C., Ghislat, G. & Ballester, P. J. A gentle introduction to understanding preclinical data for cancer pharmaco-omic modeling. Brief. Bioinform. 22, bbab312 (2021).

32. Vishwakarma, S. Machine learning models for virtual screening of molecules on cancer cell lines. (Aix-Marseille, 2019).

33. Caron, G. & Ermondi, G. Molecular Descriptors for Polarity: The Need for Going Beyond Polar Surface Area. Future Med. Chem. 8, 2013–2016 (2016).

34. Glossary | DrugBank Help Center. https://dev.drugbank.com/guides/terms.

35. Prediction of Drug-Like Properties. in Adaptive Systems in Drug Design 115–145 (CRC Press, 2002). doi:10.1201/9781498713702-10.

36. Ritchie, T. J. & Macdonald, S. J. F. The impact of aromatic ring count on compound developability – are too many aromatic rings a liability in drug design? Drug Discov. Today 14, 1011–1020 (2009).

37. Vennelakanti, V., Qi, H. W., Mehmood, R. & Kulik, H. J. When are two hydrogen bonds better than one? Accurate first-principles models explain the balance of hydrogen bond donors and acceptors found in proteins. Chem. Sci. 12, 1147–1162 (2021).

38. Sidorov, P., Naulaerts, S., Ariey-Bonnet, J., Pasquier, E. & Ballester, P. J. Predicting Synergism of Cancer Drug Combinations Using NCI-ALMANAC Data. Front. Chem. 7, 509 (2019).

39. Vishwakarma, S., Hernandez-Hernandez, S. & Ballester, P. J. Graph neural networks are promising for phenotypic virtual screening on cancer cell lines. Biol. Methods Protoc. 9, bpae065 (2024).

40. Hernández-Hernández, S., Vishwakarma, S. & Ballester, P. Conformal prediction of small-molecule drug resistance in cancer cell lines. in Proceedings of the Eleventh Symposium on Conformal and Probabilistic Prediction with Applications 92–108 (PMLR, 2022).

