## Supplementary Figure 1 for "Machine Learning-Based Drug Response Prediction Identifies Novel Therapeutic Candidates for Colorectal Cancer Cell Line KM-12"

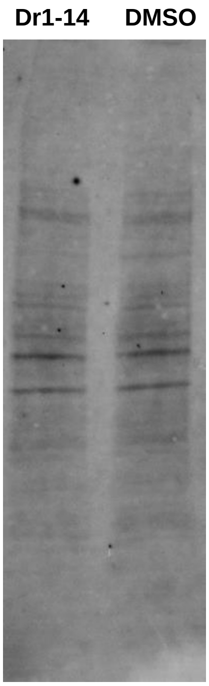


**Supplementary Figure 1.** Representative image of the total protein content using the stain free method according to the manufacturers instructions (Biorad) confirming equal loading of cell lysates generated from Dr1-14 and control treated cells for the gH2AX staining shown in Figure 3c.
