## Supplementary Table 1 for "Machine Learning-Based Drug Response Prediction Identifies Novel Therapeutic Candidates for Colorectal Cancer Cell Line KM-12"

### Supplementary Table 1. Screen 1 compounds

| Compound | Chemical name |
|---|---|
| **Dr1** | 3-(3-fluorophenyl)-N-methyl-N-[(2-methyloxolan-2-yl)methyl]-[1,2,4]triazolo[4,3-b]pyridazin-6-amine |
| **Dr2** | N-{1-oxo-1-[2-(propan-2-yl)-1,4-oxazepan-4-yl]-3-(propan-2-yloxy)propan-2-yl}acetamide |
| **Dr3** | 1-methyl-6-({[6-methyl-2-(trifluoromethyl)pyrimidin-4-yl]methyl}sulfanyl)-1H,4H,5H-pyrazolo[3,4-d]pyrimidin-4-one |
| **Dr4** | (2S)-N-[(3S,3aR,6S,6aR)-6-[(2S)-2-methoxypropanamido]-hexahydrofuro[3,2-b]furan-3-yl]-2-methoxypropanamide |
| **Dr5** | 1-(4,4-difluorooxan-3-yl)-N-{[4-(methylsulfanyl)cyclohexyl]methyl}methanesulfonamide |
| **Dr6** | 2-(dimethylamino)-3-methyl-N-[(1s,4s)-4-(2-methoxypropanamido)cyclohexyl]butanamide |
| **Dr7** | 1-cyclopropyl-N-{[1-(5-methyl-1,3,4-thiadiazole-2-carbonyl)azetidin-3-yl]methyl}-5-oxopyrrolidine-2-carboxamide |
| **Dr8** | 2-methoxy-N-[(1r,3r)-3-[(2R)-2-(dimethylamino)-3-hydroxypropanamido]cyclobutyl]pentanamide |
| **Dr9** | 2-methoxy-N-[(1s,4s)-4-[2-(2-methylpropoxy)propanamido]cyclohexyl]propanamide |
| **Dr10** | (2R)-2-(dimethylamino)-3-methyl-N-[(1r,3r)-3-[(2S)-2-methoxypropanamido]cyclobutyl]butanamide |
| **Dr11** | (2R)-2-methoxy-N-[(1s,4s)-4-(2-methoxypropanamido)cyclohexyl]propanamide |
| **Dr12** | rac-N-[(3R,4R)-1-(3-methoxypropanoyl)-4-methylpyrrolidin-3-yl]-1-(2-methylpropyl)piperidine-3-carboxamide |
| **Dr13** | rac-5-[(2R,3aS,6aS)-2-methyl-1-(oxane-3-carbonyl)-octahydropyrrolo[2,3-c]pyrrol-5-yl]-4,4-dimethyl-5-oxopentanenitrile |
| **Dr14** | rac-N-ethyl-2-(1H-imidazol-1-yl)-N-(2-{2-[(2R,5S)-5-methyloxan-2-yl]acetamido}ethyl)butanamide |
| **Dr15** | N-[(5,5-dimethylpyrrolidin-3-yl)methyl]-5,5-dimethyloxane-2-carboxamide |
| **Dr16** | 2-methoxy-3-methyl-N-[(1s,4s)-4-(2-methoxy-3-methylbutanamido)cyclohexyl]butanamide |
| **Dr17** | (2S)-2-methoxy-N-[(1s,4s)-4-[2-(2-methylpropoxy)propanamido]cyclohexyl]propanamide |
| **Dr18** | 2-methoxy-3-(methylsulfanyl)-N-[(1s,4s)-4-[(2R)-2-methoxypropanamido]cyclohexyl]propanamide |
| **Dr19** | N-(cyclopropylmethyl)-2-(3-methyloxolan-2-yl)-N-[2-({spiro[3,3]heptan-2-yl}formamido)ethyl]acetamide |
| **Dr20** | N-{2-[N-methyl-1-(2-methyloxolan-3-yl)formamido]ethyl}-2-(1H-1,2,3,4-tetrazol-1-yl)thiophene-3-carboxamide |
| **Dr21** | (2R)-2-(dimethylamino)-N-[(3RS,4SR)-4-[(2S)-2-methoxypropanamido]oxolan-3-yl]-3-methylbutanamide |
| **Dr22** | (2R)-2-(dimethylamino)-N-[(3RS,4SR)-4-[(2R)-2-methoxypropanamido]oxolan-3-yl]-3-methylbutanamide |
| **Dr23** | 2-methoxy-3-methyl-N-[(1s,4s)-4-[(2R)-2-methoxypropanamido]cyclohexyl]butanamide |
| **Dr24** | N,2-dimethyl-N-[(1-{1,3,3-trimethyl-2-oxabicyclo[2,1,1]hexane-4-carbonyl}piperidin-3-yl)methyl]propanamide |
| **Dr25** | rac-N-[(1R,2R)-2-[2-(cyclobutylmethoxy)acetamido]cyclopropyl]-2-(dimethylamino)-3-methylbutanamide |
| **Dr26** | (1R,3S)-N,1,2,2,3-pentamethyl-N-(2-{[1-(oxolan-2-yl)cyclopropyl]formamido}ethyl)cyclopentane-1-carboxamide |
| **Dr27** | N-(1-{[1-(5-ethyl-5-methyloxolan-2-yl)-N-methylformamido]methyl}cyclopropyl)-5-methyl-2-oxo-2,3-dihydro-1H-imidazole-4-carboxamide |
| **Dr28** | 4-methyl-2-[(1-{3-[(2-methylcyclopropyl)formamido]propanoyl}piperidin-4-yl)methyl]-1,3-thiazole-5-carboxylic acid |
| **Dr29** | 2-cyclopropyl-N-(1-{[1-(1,1-difluoroethyl)cyclopropyl]formamido}-4-methoxybutan-2-yl)-2-methylpropanamide |
| **Dr30** | 5-(1,1-dimethyl-1,3-dihydro-2-benzofuran-5-yl)-1,3-benzoxazol-2-amine |
| **Dr31** | 5-[3-(oxan-4-yl)-4,5-dihydro-1,2-oxazol-5-yl]-3-{[1-(2,2,2-trifluoroethyl)-1H-pyrazol-3-yl]methyl}-1,2,4-oxadiazole |
| **Dr32** | 5,8-difluoro-N-({5H,7H,8H-pyrano[4,3-d]pyrimidin-2-yl}methyl)quinolin-2-amine |
| **Dr33** | rac-3-[4-({[(3aR,6aR)-hexahydro-2H-cyclopenta[b]furan-6a-yl]methyl}amino)-5-chloro-6-oxo-1,6-dihydropyridazin-1-yl]-1??-thiolane-1,1-dione |
| **Dr34** | 1-methyl-6-({[4-methyl-5-(thiophen-2-yl)-4H-1,2,4-triazol-3-yl]methyl}sulfanyl)-1H,4H,5H-pyrazolo[3,4-d]pyrimidin-4-one |
| **Dr35** | 5-{5-[3-(3,6-dihydro-2H-pyran-4-yl)phenyl]-1,2,4-oxadiazol-3-yl}-3-methoxy-2-methylpyridine |
| **Dr36** | (2R)-2-(dimethylamino)-3-methyl-N-[(1s,3s)-3-(2-methoxypropanamido)cyclobutyl]butanamide |
| **Dr37** | 2-methoxy-3-methyl-N-[(1s,4s)-4-[2-(2-methylpropoxy)propanamido]cyclohexyl]butanamide |
| **Dr38** | (2R)-2-methoxy-N-[(1s,4s)-4-[2-(2-methylpropoxy)propanamido]cyclohexyl]propanamide |
| **Dr39** | 2-(dimethylamino)-3-methyl-N-[(1s,3s)-3-[(2S)-2-methoxypropanamido]cyclobutyl]butanamide |
| **Dr40** | 2-(1-{6-azaspiro[3,5]nonane-8-carbonyl}piperidin-4-yl)acetonitrile |
| **Dr41** | (2R)-2-(dimethylamino)-3-methyl-N-[(1s,3s)-3-[2-(2-methylpropoxy)propanamido]cyclobutyl]butanamide |
| **Dr42** | 2-methoxy-3-methyl-N-[(1s,4s)-4-(2-methoxypropanamido)cyclohexyl]butanamide |
| **Dr43** | N2-[(1RS,2RS)-2-[(2S)-2-(dimethylamino)-3-methylbutanamido]cyclopropyl]thiophene-2,5-dicarboxamide |
| **Dr44** | (2S)-2-methoxy-N-[(1s,4s)-4-[(2R)-2-methoxypropanamido]cyclohexyl]propanamide |
| **Dr45** | (2S)-2-methoxy-N-[(1s,4s)-4-[(2S)-2-methoxypropanamido]cyclohexyl]propanamide |
| **Dr46** | 4-methyl-N-[(1r,3r)-3-[(2R)-2-[2-(dimethylamino)ethoxy]propanamido]cyclobutyl]morpholine-3-carboxamide |
| **Dr47** | (2R)-N-[(3S,3aR,6S,6aR)-6-[(2R)-2-methoxypropanamido]-hexahydrofuro[3,2-b]furan-3-yl]-2-(dimethylamino)-3-methylbutanamide |
| **Dr48** | N-[(3S,3aR,6S,6aR)-6-[(2R)-2-methoxypropanamido]-hexahydrofuro[3,2-b]furan-3-yl]-2-(dimethylamino)-3-methylbutanamide |
| **Dr49** | (2S)-N-[(3S,3aR,6S,6aR)-6-[(2S)-2-methoxypropanamido]-hexahydrofuro[3,2-b]furan-3-yl]-2-(dimethylamino)-3-methylbutanamide |
| **Dr50** | (2R)-N-[(3S,3aR,6S,6aR)-6-(2-methoxypropanamido)-hexahydrofuro[3,2-b]furan-3-yl]-2-(dimethylamino)-3-methylbutanamide |
| **Dr51** | (2S)-N-[(3S,3aR,6S,6aR)-6-[2-(2-methylpropoxy)propanamido]-hexahydrofuro[3,2-b]furan-3-yl]-2-methoxypropanamide |
| **Dr52** | N-[(3S,3aR,6S,6aR)-6-[(2S)-2-methoxypropanamido]-hexahydrofuro[3,2-b]furan-3-yl]-2-methoxy-3-methylbutanamide |
| **Dr53** | N-[(3S,3aR,6S,6aR)-6-[(2S)-2-methoxypropanamido]-hexahydrofuro[3,2-b]furan-3-yl]-2-(dimethylamino)-3-methylbutanamide |
| **Dr54** | 5-{1-ethyl-5-[1-(2-methylphenyl)-1H-pyrazol-3-yl]-1H-1,2,4-triazol-3-yl}-2-methoxypyridine |
| **Dr55** | 3-methoxy-2-methyl-5-{5-[5-(oxan-4-yl)-1,2-oxazol-3-yl]-1,2,4-oxadiazol-3-yl}pyridine |
| **Dr56** | N-(cyclopropylmethyl)-N-{2-[(3,3-dimethyloxetan-2-yl)formamido]ethyl}thiane-2-carboxamide |
| **Dr57** | (2R)-N-[(3S,3aR,6S,6aR)-6-[(2S)-2-methoxypropanamido]-hexahydrofuro[3,2-b]furan-3-yl]-2-(dimethylamino)-3-methylbutanamide |
| **Dr58** | N-[(3S,3aR,6S,6aR)-6-[(2R)-2-methoxypropanamido]-hexahydrofuro[3,2-b]furan-3-yl]-2-methoxypropanamide |
| **Dr59** | N-[(3S,3aR,6S,6aR)-6-[(2R)-2-methoxypropanamido]-hexahydrofuro[3,2-b]furan-3-yl]-2-methoxy-3-methylbutanamide |
| **Dr60** | (2R)-N-[(3S,3aR,6S,6aR)-6-[2-(2-methylpropoxy)propanamido]-hexahydrofuro[3,2-b]furan-3-yl]-2-methoxypropanamide |
| **Dr61** | N-[(3S,3aR,6S,6aR)-6-[(2R)-2-methoxypropanamido]-hexahydrofuro[3,2-b]furan-3-yl]-2-methoxy-3-(methylsulfanyl)propanamide |
| **Dr62** | N-[(3S,3aR,6S,6aR)-6-[(2R)-2-methoxypropanamido]-hexahydrofuro[3,2-b]furan-3-yl]-2-methoxybutanamide |
| **Dr63** | rac-4-{[(2R,3S)-3-methyloxolan-2-yl]methyl}-2-(propan-2-yl)thiomorpholine |
| **Dr64** | rac-N-[(3R,4R)-1-{2-[carbamoyl(methyl)amino]acetyl}-4-methylpyrrolidin-3-yl]-3-methanesulfonyl-2-methylpropanamide |
| **Dr65** | (2S)-N-[(2S)-1,1-difluoropropan-2-yl]-2-(dimethylamino)-3-methylbutanamide |
| **Dr66** | N-[(4-tert-butylcyclohexyl)methyl]-1-{2-oxabicyclo[2,2,2]octan-4-yl}methanesulfonamide |
| **Dr67** | (2R)-N-[(3S,3aR,6S,6aR)-6-[(2R)-2-methoxypropanamido]-hexahydrofuro[3,2-b]furan-3-yl]-2-methoxypropanamide |
| **Dr68** | 1-[6-(aminomethyl)spiro[3,3]heptane-2-carbonyl]-N-[2-(dimethylamino)ethyl]piperidine-4-carboxamide |
| **Dr69** | rac-N2-[(1R,2R)-2-[2-(dimethylamino)-3-methylbutanamido]cyclopropyl]thiophene-2,5-dicarboxamide |
| **Dr70** | rac-2,2-dimethyl-3-[({[(2R,3S)-3-methyloxolan-2-yl]methyl}sulfanyl)methyl]-1,4-dioxane |
| **Dr71** | N-[(3S,3aR,6S,6aR)-6-[(2R)-2-methoxypropanamido]-hexahydrofuro[3,2-b]furan-3-yl]-2,3-dimethoxypropanamide |
| **Dr72** | rac-2-(4-ethylmorpholin-2-yl)ethyl (2R,3R)-1-ethyl-6-oxo-2-(pyridin-4-yl)piperidine-3-carboxylate |
| **Dr73** | (2S)-2-(dimethylamino)-N-[(1RS,2RS)-2-[(2S)-2-(dimethylamino)propanamido]cyclopropyl]-3-hydroxypropanamide |
| **Dr74** | 1-{1,8-dioxaspiro[4,5]decan-2-yl}-N-[(1r,4r)-4-cyanocyclohexyl]methanesulfonamide |
