## Supplementary Table 2 for "Machine Learning-Based Drug Response Prediction Identifies Novel Therapeutic Candidates for Colorectal Cancer Cell Line KM-12"

### Supplementary Table 2. Screen 2 compounds

| Compound | Chemical name |
|---|---|
| **Dr1-1** | 4-{[(E)-{3-chloro-5-ethoxy-4-[(3-fluorobenzyl)oxy]phenyl}methylidene]amino}-5-phenyl-4H-1,2,4-triazole-3-thiol |
| **Dr1-2** | 6-(2,6-dimethoxyphenoxy)-3-phenyl[1,2,4]triazolo[4,3-b]pyridazine |
| **Dr1-3** | N-(3-methylbutyl)-3-phenyl[1,2,4]triazolo[4,3-b]pyridazin-6-amine |
| **Dr1-4** | N-[2-(cyclohex-1-en-1-yl)ethyl]-1-(3-phenyl[1,2,4]triazolo[4,3-b]pyridazin-6-yl)piperidine-4-carboxamide |
| **Dr1-5** | (2Z)-2-{[5-(3-fluorophenyl)furan-2-yl]methylidene}-6,7-dimethyl[1,3]thiazolo[3,2-a]benzimidazol-3(2H)-one |
| **Dr1-6** | 3-(4-bromophenyl)-6-(2,6-dimethoxyphenoxy)[1,2,4]triazolo[4,3-b]pyridazine |
| **Dr1-7** | 3-[3-(benzyloxy)phenyl]-6-(trifluoromethyl)[1,2,4]triazolo[4,3-a]pyridine |
| **Dr1-8** | N-[(3-ethoxyoxolan-3-yl)methyl]-N-methyl-3-(thiophen-3-yl)-[1,2,4]triazolo[4,3-b]pyridazin-6-amine |
| **Dr1-9** | 1-methyl-5-({3-phenyl-[1,2,4]triazolo[4,3-b]pyridazin-6-yl}sulfanyl)-1H-1,2,4-triazole |
| **Dr1-10** | [(2S,4S)-4-fluoro-1-{3-phenyl-[1,2,4]triazolo[4,3-b]pyridazin-6-yl}pyrrolidin-2-yl]methanol |
| **Dr1-11** | 1-(2-fluoroethyl)-N-[3-(4-fluorophenyl)-[1,2,4]triazolo[4,3-b]pyridazin-6-yl]-1H-pyrazol-4-amine |
| **Dr1-12** | 3-phenyl-N-({5H,6H,7H,8H-[1,2,4]triazolo[4,3-a]pyridin-3-yl}methyl)-[1,2,4]triazolo[4,3-b]pyridazin-6-amine |
| **Dr1-13** | 3-(3-methoxyphenyl)-4-methyl-5-[(2-phenoxyethyl)sulfanyl]-4H-1,2,4-triazole |
| **Dr1-14** | N-[(1-benzyl-1H-pyrazol-4-yl)methyl]-N-methyl-3-phenyl-[1,2,4]triazolo[4,3-b]pyridazin-6-amine |
| **Dr1-15** | 3-[(1R)-1-({3-phenyl-[1,2,4]triazolo[4,3-b]pyridazin-6-yl}amino)ethyl]phenol |
| **Dr1-16** | N-[2-(1-methyl-1H-imidazol-5-yl)ethyl]-3-phenyl-[1,2,4]triazolo[4,3-b]pyridazin-6-amine |
| **Dr1-17** | 3-methyl-2-phenyl-1-{3-phenyl-[1,2,4]triazolo[4,3-b]pyridazin-6-yl}azetidine |
| **Dr1-18** | 3,5-dimethyl-1-(1-{3-phenyl-[1,2,4]triazolo[4,3-b]pyridazin-6-yl}azetidin-3-yl)-1H-pyrazole |
| **Dr1-19** | N-[(3,5-dimethyl-1,2-oxazol-4-yl)methyl]-N-methyl-3-phenyl-[1,2,4]triazolo[4,3-b]pyridazin-6-amine |
| **Dr1-20** | N-[(4-ethylmorpholin-2-yl)methyl]-3-phenyl-[1,2,4]triazolo[4,3-b]pyridazin-6-amine |
| **Dr1-21** | 3-ethyl-5-({3-phenyl-[1,2,4]triazolo[4,3-b]pyridazin-6-yl}sulfanyl)-4H-1,2,4-triazole |
| **Dr1-22** | 5-{3-phenyl-[1,2,4]triazolo[4,3-b]pyridazin-6-yl}-2-thia-5-azabicyclo[4.2.0]octane |
| **Dr1-23** | 6-[(4aS,7aR)-4-methyl-octahydropyrrolo[3,4-b][1,4]oxazin-6-yl]-3-phenyl-[1,2,4]triazolo[4,3-b]pyridazine |
| **Dr1-24** | (3R,5S)-1-[3-(3-fluorophenyl)-[1,2,4]triazolo[4,3-b]pyridazin-6-yl]-5-(methoxymethyl)pyrrolidin-3-ol |
| **Dr1-25** | N-[(3-methoxyoxolan-3-yl)methyl]-N-methyl-3-(thiophen-3-yl)-[1,2,4]triazolo[4,3-b]pyridazin-6-amine |
| **Dr1-26** | 3-(5-{1-[(2-methyltetrahydrofuran-2-yl)methyl]-1H-imidazol-2-yl}-2-furyl)-1H-pyrazole |
| **Dr1-27** | 6-(3-fluorophenyl)-N-methyl-N-[2-(tetrahydro-2H-pyran-4-yl)ethyl]imidazo[2,1-b][1,3]thiazole-3-carboxamide |
| **Dr1-28** | 6-(3-fluorophenyl)-N-[2-(1H-imidazol-1-yl)ethyl]-N-methylimidazo[2,1-b][1,3]thiazole-3-carboxamide |
| **Dr1-29** | 6-(3-fluorophenyl)-N-(3-hydroxy-3-methylbutyl)-N-methylimidazo[2,1-b][1,3]thiazole-3-carboxamide |
| **Dr1-30** | 2-(3-[1,2,4]triazolo[4,3-a]pyridin-3-ylphenoxy)ethanol |
| **Dr1-31** | 1-({cyclopropyl[(3-fluorophenyl)methyl]amino}methyl)-4-ethyl-3-(furan-2-yl)-4,5-dihydro-1H-1,2,4-triazole-5-thione |
| **Dr1-32** | [(3-oxo-1-phenylbutan-2-yl)carbamoyl]methyl 3-[3-(4-fluorophenyl)-1-phenyl-1H-pyrazol-4-yl]prop-2-enoate |
| **Dr1-33** | 3-{[2-(2-fluorophenoxy)ethyl]sulfanyl}-5-(3-methoxyphenyl)-4-phenyl-4H-1,2,4-triazole |
| **Dr1-34** | ethyl 2-{[3-(3,5-dichlorophenyl)-[1,2,4]triazolo[4,3-b]pyridazin-6-yl](methyl)amino}acetate |
| **Dr1-35** | 3-nitro-1-{3-phenyl-[1,2,4]triazolo[4,3-b]pyridazin-6-yl}-1,4-dihydropyridin-4-one |
| **Dr1-36** | 6-{3-ethyl-5H,6H,7H,8H-[1,2,4]triazolo[4,3-a]pyrazin-7-yl}-3-phenyl-[1,2,4]triazolo[4,3-b]pyridazine |
| **Dr1-37** | N-methyl-N-(2-phenoxyethyl)-3-phenyl-[1,2,4]triazolo[4,3-b]pyridazin-6-amine |
| **Dr1-38** | N-methyl-N-[2-(2-methylphenoxy)ethyl]-3-(trifluoromethyl)-[1,2,4]triazolo[4,3-b]pyridazin-6-amine |
| **Dr1-39** | 4-ethyl-3-(furan-2-yl)-1-{[methyl({[4-(trifluoromethyl)phenyl]methyl})amino]methyl}-4,5-dihydro-1H-1,2,4-triazole-5-thione |
