## Supplementary Table 3 for "Machine Learning-Based Drug Response Prediction Identifies Novel Therapeutic Candidates for Colorectal Cancer Cell Line KM-12"

### Supplementary Table 3. Screen 3 compounds

| Compound | Chemical name |
|---|---|
| **Dr1-40** | N-methyl-N-[(oxolan-2-yl)methyl]-3-(pyridin-4-yl)-[1,2,4]triazolo[4,3-b]pyridazin-6-amine |
| **Dr1-41** | N-methyl-N-[(2-methyloxolan-2-yl)methyl]-3-(pyridin-4-yl)-[1,2,4]triazolo[4,3-b]pyridazin-6-amine |
| **Dr1-42** | 4,4-difluoro-1-[3-(3-fluorophenyl)-[1,2,4]triazolo[4,3-b]pyridazin-6-yl]-2-methylpyrrolidine |
| **Dr1-43** | 3-(3-fluorophenyl)-N-[(1-methoxycyclopentyl)methyl]-N-methyl-[1,2,4]triazolo[4,3-b]pyridazin-6-amine |
| **Dr1-44** | N-methyl-N-[(2-methyloxolan-2-yl)methyl]-3-(pyridin-2-yl)-[1,2,4]triazolo[4,3-b]pyridazin-6-amine |
| **Dr1-45** | N'-[1-(2-fluoropyridin-4-yl)-1H-pyrazol-4-yl]-N-methyl-N-[(2-methyloxolan-2-yl)methyl]ethanediamide |
| **Dr1-46** | 2-methyl-3-[methyl({3-phenyl-[1,2,4]triazolo[4,3-b]pyridazin-6-yl})amino]propanenitrile |
| **Dr1-47** | N-methyl-N-[(2-methyloxolan-2-yl)methyl]-3-phenyl-[1,2,4]triazolo[4,3-b]pyridazin-6-amine |
| **Dr1-48** | N-methyl-N-[(2-methyloxolan-2-yl)methyl]-3-(pyridin-3-yl)-[1,2,4]triazolo[4,3-b]pyridazin-6-amine |
| **Dr1-49** | 4-{[3-(3-fluorophenyl)-[1,2,4]triazolo[4,3-b]pyridazin-6-yl](methyl)amino}-3,3-dimethylbutan-1-ol |
| **Dr1-50** | N-[(1-fluorocyclopentyl)methyl]-3-(3-fluorophenyl)-N-methyl-[1,2,4]triazolo[4,3-b]pyridazin-6-amine |
| **Dr1-51** | 3-(3-fluorophenyl)-N-methyl-N-[2-(prop-2-yn-1-yloxy)ethyl]-[1,2,4]triazolo[4,3-b]pyridazin-6-amine |
| **Dr1-52** | 3-(3-fluorophenyl)-N-methyl-N-[(1H-pyrazol-4-yl)methyl]-[1,2,4]triazolo[4,3-b]pyridazin-6-amine |
| **Dr1-53** | 1-[3-(3-fluorophenyl)-[1,2,4]triazolo[4,3-b]pyridazin-6-yl]-2,3-dimethylpiperidine |
| **Dr1-54** | N-methyl-3-(2-methylcyclopropyl)-N-[(2-methyloxolan-2-yl)methyl]-[1,2,4]triazolo[4,3-b]pyridazin-6-amine |
| **Dr1-55** | N-[(cyclopent-1-en-1-yl)methyl]-3-(3-fluorophenyl)-[1,2,4]triazolo[4,3-b]pyridazin-6-amine |
| **Dr1-56** | 2-{[3-(3-fluorophenyl)-[1,2,4]triazolo[4,3-b]pyridazin-6-yl](methyl)amino}-N-methylacetamide |
| **Dr1-57** | 3-(3-fluorophenyl)-N-[(2-methyloxolan-2-yl)methyl]-[1,2,4]triazolo[4,3-b]pyridazin-6-amine |
| **Dr1-58** | N-[(1-fluorocyclobutyl)methyl]-3-(3-fluorophenyl)-N-methyl-[1,2,4]triazolo[4,3-b]pyridazin-6-amine |
| **Dr1-59** | 3-(3-fluorophenyl)-N-methyl-N-{[(2R)-oxolan-2-yl]methyl}-[1,2,4]triazolo[4,3-b]pyridazin-6-amine |
| **Dr1-60** | 3-(3-fluorophenyl)-N-[(3-methoxyoxolan-3-yl)methyl]-N-methyl-[1,2,4]triazolo[4,3-b]pyridazin-6-amine |
| **Dr1-61** | N-[3-(dimethylamino)propyl]-3-(3-fluorophenyl)-N-methyl-[1,2,4]triazolo[4,3-b]pyridazin-6-amine |
| **Dr1-62** | 2-methyl-2-({methyl[3-(pyridin-4-yl)-[1,2,4]triazolo[4,3-b]pyridazin-6-yl]amino}methyl)propane-1,3-diol |
| **Dr1-63** | 3-{[3-(3-fluorophenyl)-[1,2,4]triazolo[4,3-b]pyridazin-6-yl](methyl)amino}propanenitrile |
| **Dr1-64** | 3-{[3-(3-fluorophenyl)-[1,2,4]triazolo[4,3-b]pyridazin-6-yl](methyl)amino}-2-methylpropanenitrile |
| **Dr1-65** | N-[(3-fluorocyclobutyl)methyl]-3-(3-fluorophenyl)-N-methyl-[1,2,4]triazolo[4,3-b]pyridazin-6-amine |
| **Dr1-66** | N-[(2,2-difluorocyclopropyl)methyl]-3-(3-fluorophenyl)-N-methyl-[1,2,4]triazolo[4,3-b]pyridazin-6-amine |
| **Dr1-67** | 2-({[3-(3-fluorophenyl)-[1,2,4]triazolo[4,3-b]pyridazin-6-yl](methyl)amino}methyl)-2-methylpropane-1,3-diol |
| **Dr1-68** | 3-{[3-(3-fluorophenyl)-[1,2,4]triazolo[4,3-b]pyridazin-6-yl](methyl)amino}-2,2-dimethylpropan-1-ol |
| **Dr1-69** | 7-[3-(3-fluorophenyl)-[1,2,4]triazolo[4,3-b]pyridazin-6-yl]-1,4-dioxa-7-azaspiro[4.4]nonane |
| **Dr1-70** | 1-[3-(3-fluorophenyl)-[1,2,4]triazolo[4,3-b]pyridazin-6-yl]-2-methylazepane |
| **Dr1-71** | N-[3-(dimethylamino)-2,2-dimethylpropyl]-3-(3-fluorophenyl)-N-methyl-[1,2,4]triazolo[4,3-b]pyridazin-6-amine |
| **Dr1-72** | 2-(2-{[3-(3-fluorophenyl)-[1,2,4]triazolo[4,3-b]pyridazin-6-yl](methyl)amino}ethoxy)ethan-1-ol |
| **Dr1-73** | 1-[3-(3-fluorophenyl)-[1,2,4]triazolo[4,3-b]pyridazin-6-yl]-2,4-dimethylpiperazine |
| **Dr1-74** | 3-(3-fluorophenyl)-N-methyl-N-(3-methylcyclopentyl)-[1,2,4]triazolo[4,3-b]pyridazin-6-amine |
| **Dr1-75** | N-methyl-N-{[(2R)-oxolan-2-yl]methyl}-3-(pyridin-4-yl)-[1,2,4]triazolo[4,3-b]pyridazin-6-amine |
| **Dr1-76** | N-(but-3-en-1-yl)-3-(3-fluorophenyl)-N-methyl-[1,2,4]triazolo[4,3-b]pyridazin-6-amine |
| **Dr1-77** | N-methyl-N-[(2-methyloxolan-2-yl)methyl]-3-[4-(5-methylpyridin-3-yl)-1H-1,2,3-triazol-1-yl]propanamide |
| **Dr1-78** | 3-{[3-(4-fluorophenyl)-[1,2,4]triazolo[4,3-b]pyridazin-6-yl](methyl)amino}-2-methylpropanenitrile |
| **Dr1-79** | 3-(3-fluorophenyl)-N-[(furan-2-yl)methyl]-N-methyl-[1,2,4]triazolo[4,3-b]pyridazin-6-amine |
| **Dr1-80** | 3-(3-fluorophenyl)-N-methyl-N-{[(2R)-pyrrolidin-2-yl]methyl}-[1,2,4]triazolo[4,3-b]pyridazin-6-amine |
| **Dr1-81** | 3-(3-fluorophenyl)-N-methyl-N-[(thiolan-2-yl)methyl]-[1,2,4]triazolo[4,3-b]pyridazin-6-amine |
| **Dr1-82** | 3-(3-fluorophenyl)-N-methyl-N-[2-(methylamino)ethyl]-[1,2,4]triazolo[4,3-b]pyridazin-6-amine |
| **Dr1-83** | N-[2-(2-aminoethoxy)ethyl]-3-(3-fluorophenyl)-N-methyl-[1,2,4]triazolo[4,3-b]pyridazin-6-amine |
| **Dr1-84** | 3-methyl-3-[(2-methyloxolan-2-yl)methyl]-1-[(5-phenyl-1H-pyrazol-3-yl)methyl]urea |
| **Dr1-85** | (2S)-1-[3-(3-fluorophenyl)-[1,2,4]triazolo[4,3-b]pyridazin-6-yl]-2-methylpiperazine |
| **Dr1-86** | (2R)-1-[3-(3-fluorophenyl)-[1,2,4]triazolo[4,3-b]pyridazin-6-yl]-2-methylpiperazine |
| **Dr1-87** | 3-{[1-(3-methoxyphenyl)pyrrolidin-3-yl]methyl}-1-methyl-1-[(2-methyloxolan-2-yl)methyl]urea |
| **Dr1-88** | ethyl 2-{[3-(3-fluorophenyl)-[1,2,4]triazolo[4,3-b]pyridazin-6-yl](methyl)amino}acetate |
| **Dr1-89** | N,3-dimethyl-N-[(2-methyloxolan-2-yl)methyl]-[1,2,4]triazolo[4,3-b]pyridazin-6-amine |
| **Dr1-90** | 1-({[1-(3-chlorophenyl)-1H-pyrazol-3-yl]methyl}amino)-3-(2-methyloxolan-2-yl)propan-2-ol |
| **Dr1-91** | 3-(3-fluorophenyl)-N-methyl-N-[(oxolan-2-yl)methyl]-[1,2,4]triazolo[4,3-b]pyridazin-6-amine |
| **Dr1-92** | 3-(3-fluorophenyl)-N-methyl-N-[(pyrrolidin-2-yl)methyl]-[1,2,4]triazolo[4,3-b]pyridazin-6-amine |
| **Dr1-93** | N-(2-aminopropyl)-3-(3-fluorophenyl)-N-methyl-[1,2,4]triazolo[4,3-b]pyridazin-6-amine |
| **Dr1-94** | 1-[3-(3-fluorophenyl)-[1,2,4]triazolo[4,3-b]pyridazin-6-yl]-2-methylpiperazine |
| **Dr1-95** | 3-(3-fluorophenyl)-N-methyl-N-{[(2S)-pyrrolidin-2-yl]methyl}-[1,2,4]triazolo[4,3-b]pyridazin-6-amine |
| **Dr1-96** | N-[2-(dimethylamino)ethyl]-3-(3-fluorophenyl)-N-methyl-[1,2,4]triazolo[4,3-b]pyridazin-6-amine |
| **Dr1-97** | N-methyl-3-phenyl-N-[(thiolan-2-yl)methyl]-[1,2,4]triazolo[4,3-b]pyridazin-6-amine |
| **Dr1-98** | 3-(3-fluorophenyl)-N-methyl-N-[(3-methyloxetan-3-yl)methyl]-[1,2,4]triazolo[4,3-b]pyridazin-6-amine |
| **Dr1-99** | N-(2-amino-2-methylpropyl)-3-(3-fluorophenyl)-N-methyl-[1,2,4]triazolo[4,3-b]pyridazin-6-amine |
| **Dr1-100** | 1-(3-fluorophenyl)-N-[2-methyl-3-(2-methyl-1,3-dioxolan-2-yl)propyl]-2-oxopyrrolidine-3-carboxamide |
| **Dr1-101** | 3-{[3-(3-fluorophenyl)-[1,2,4]triazolo[4,3-b]pyridazin-6-yl]amino}pentan-2-ol; trifluoroacetic acid |
| **Dr1-102** | N-methyl-3-(pyridin-3-yl)-N-[(thiolan-2-yl)methyl]-[1,2,4]triazolo[4,3-b]pyridazin-6-amine |
