## Supplementary Table 4 for "Machine Learning-Based Drug Response Prediction Identifies Novel Therapeutic Candidates for Colorectal Cancer Cell Line KM-12"

**Supplementary Table 4. Efficacy of tested compounds (2.5 μM) to decrease cell viability of KM-12 cells below 90 % in 72 h assays.**

| Screen | Tested compounds | Compounds with target efficacy (n) | Compounds with target efficacy (%) |
|---|---|---|---|
| Screen 1 | 74 | 1 | 1.4 |
| Screen 2 | 39 | 3 | 7.7 |
| Screen 3 | 63 | 22 | 35 |
